# The Conformation of the Complementary Strand and the Deformation of the DNA Groove upon DDB2 Binding Justifies the Different Repair Rates for Cyclobutane Pyrimidine Dimers

**DOI:** 10.64898/2026.05.10.724087

**Authors:** Yani Kedjar, Cécilia Hognon, Thierry Douki, Elise Dumont, Antonio Monari

## Abstract

The repair of photo-induced DNA lesions through nucleotide excision repair machinery is still the source of important questions. It has been observed that the repair rate of the different cyclobutane pyrimidine dimers, i.e. the photoproducts induced by dimerization of two π-stacked pyrimidines (T<>T, T<>C, C<>T, C<>C), depends on the nucleobases involved in the lesion. TT derivatives (T<>T) are removed more slowly than those containing cytosine, especially in 5’. Using all-atom molecular dynamics simulations, we demonstrate that the variation of the repair rate observed in human skin and in cultured cutaneous cell may be associated to the recognition of the four lesions by the DDB2 protein moiety, and more specifically by the differential structural deformation induced on the complementary strand and the major groove. These effects may then hamper differentially the downstream recruitment of the repair complexes. The observed DNA deformation correlates with the experimental repair rate and suggests a structural rationale for the different repair rates of CPD by nucleotide excision repair machinery.

**GRAPHICAL ABSTRACT:** 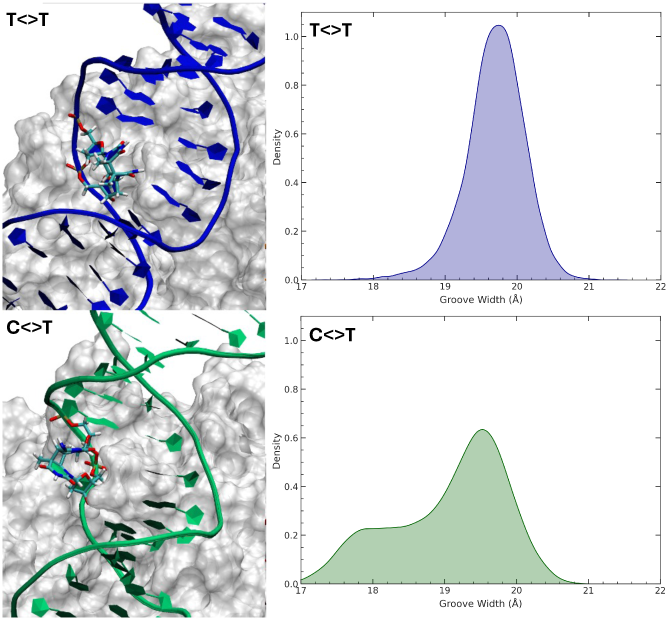

## INTRODUCTION

Overexposure to sunlight is recognized as a cause of the appearance of malignant skin lesions that may evolve to skin cancer.^1–8^ It is well established that, despite the global photostability of DNA,^9^ UV light can induce the formation of DNA photolesions, mostly in the form of cyclobutane pyrimidine dimers (CPD, see Figure 1A), upon direct absorption of UVB light by the DNA nucleobases.^10^ CPDs are formed at the four possible sequences harboring two adjacent pyrimidines, namely CC, CT, TC and TT.^3,4,11–13^ UVB radiation also triggers the formation of pyrimidine (6-4) pyrimidone photoproducts (64-PP) present in a threefold lower yield than CPDs.^11,14–17^ These two lesions are harmful for the cell^18^ for two antinomic reasons. Indeed, while CPDs are characterized by a low repair rate, leading to accumulation of these defects in cells, 64-PPs, however extremely mutagenic, are much faster repaired than CPDs.^1^ The differential effects have been correlated to the related change produced in the DNA mechanical properties, and to flexibility.^19–23^ CPDs tend to rigidify the whole oligonucleotide with minor structural differences, compared to native B-DNA, whereas 64-PPs exhibit an extended polymorphism characterized by the simultaneous presence of very different conformations, and their rapid interconversion over the microsecond timescale.^24^ Hence, while CPDs may efficiently escape recognition by the repair machinery, namely nucleotide excision repair (NER), 64-PPs are easily recognized, but their flexibility induces a much lesser replication resistance or fork blockages.^25–28^ Important differences in the processing rate of the four CPDs have been observed based on highly specific mass spectrometry measurements.^29^ The quantification of their repair rate in human skin cells, either fibroblasts or keratinocytes, indicates that the processing efficiency can greatly vary along the four possible adducts, as shown in Figure 1B.

**Figure 1.**
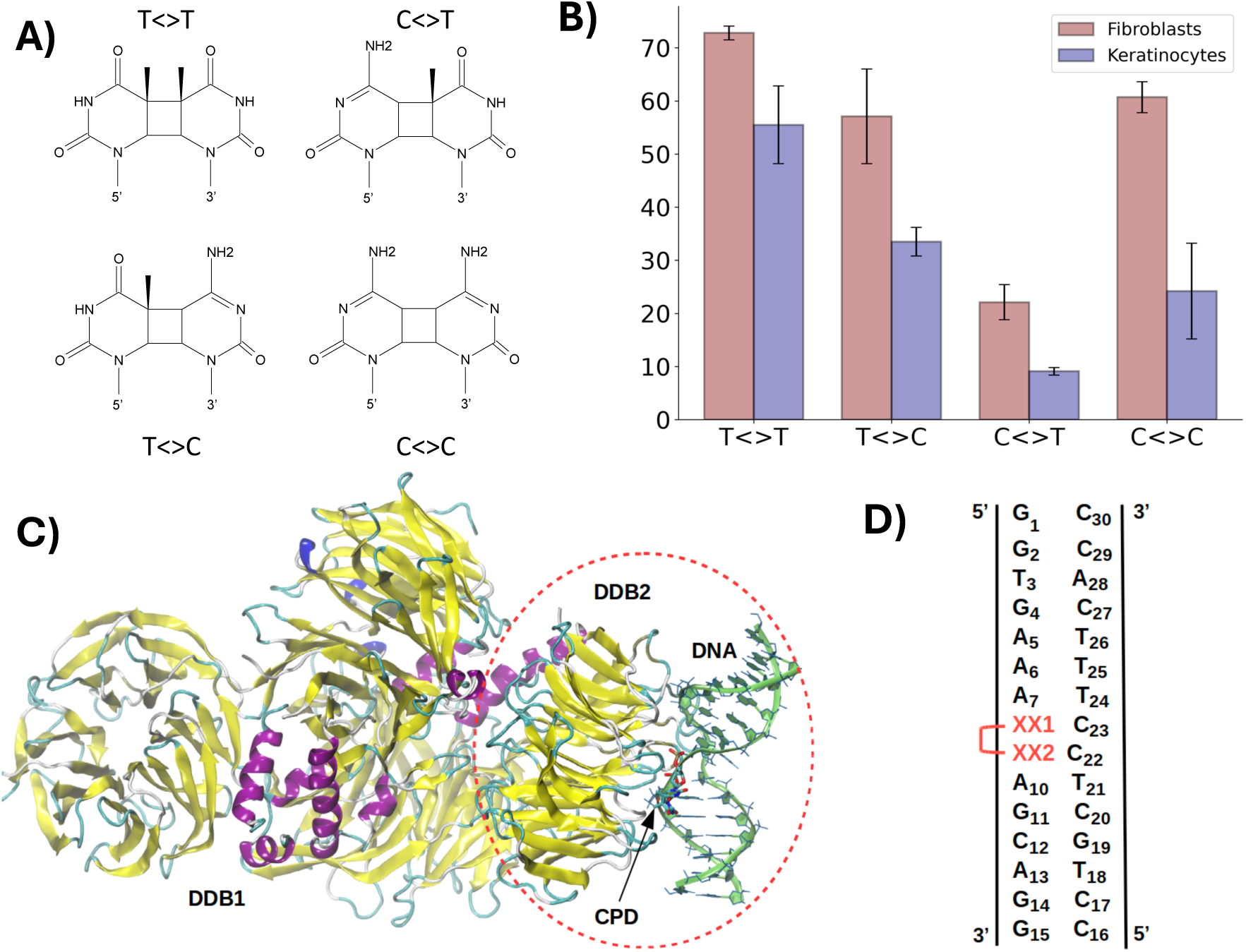
A) The chemical structures of the specific CPD investigated here, i.e. T<>T, C<>T, T<>C, C<>C. B) Repair with respect to CPD lesions and expressed as percentage of lesion still present after 24h from light exposure. Data adapted from reference^29^ error bars are indicated. C) Crystal structure of DDB1-DDB2 protein bound to a CPD (T<>T) containing duplex, PDB: 4A09.^30^ We only simulated the DNA recognition subunits (DDB2) evidenced with red dashed lines. D) Schematization of the DNA sequence containing the CPD lesion (marked as XX).

T<>T undergoes a slow repair and may persist up to a level of 70%, in stark contrast with C<>T that may reach levels of less than 10%. T<>C and C<>C exhibit intermediate repair rates. From the analysis of the data reported in Figure 1B, it is, therefore, possible to sketch out a repair-efficiency performance order, which can be summarized as follows:

“T<>T” < “T<>C” << “C<>C” < “C<>T”

Hence, it is apparent that the presence of a cytosine at the 5’ position results in a net increase of the repair efficiency. This order of repair rate is reminiscent of the mutation frequency in skin tumors and UV-irradiated cells.^31,32^ Indeed, the most frequent mutations are observed at the 3’-end of T<>C sequences, and, to a lesser extent, at C<>C sites, while C<>T sequences are very rarely mutagenic.^29^ Mutations at T<>T sites are also quite rare, but this is explained by the fact that the two thymines in a T<>T CPD preserve their coding properties upon replication.^33–35^

To explain such differences in the repair rate, we turn to molecular dynamics (MD) simulations, allowing molecular biological processes to be described at the atomic level.^36–41^ Specifically, we compare the behavior of oligonucleotides containing the four different CPD to characterize possible inherent structural signatures associated to the different repair rates. Subsequently, the interaction of a damaged DNA strand with the nuclear excision repair^42,43^ (NER) protein complex DDB1/DDB2,^44–48^ responsible for the recognition of photolesions and the triggering of the repair process, is also considered to unravel the molecular bases underlying the different repair rates. Indeed, upon interaction with the DDB1/DDB2 complex the lesioned nucleobases are extruded from the double helical arrangements. The extrusion of the damaged bases is then translated into a pronounced structural modification of the complementary strand, particularly at -1 (3’) and +1 (5’) position from the lesion, which facilitates recognition by the subsequent NER enzymes, notably the XPC-RAD23B complex.^49^ The binding of XPC-RAD23B triggers the recruitment of ubiquitination agents which induces a topological transition towards less compact chromatin and hence favors the accessibility of the other repair enzymes.

To model the first events of the repair machinery, we have considered the complex crystallized by Scrima et al. (PDB code 4a09),^30^ depicted in Figure 1C. The crystal structure consists of the DDB1/DDB2 complex bound to a 15-bp oligonucleotide containing a T<>T lesion. In what follows, we focused on the interaction between DDB2 and DNA, while the DDB1 moiety was removed. This choice is justified by the low resolution of the DDB1 unit,^30^ due do its greater flexibility, and by the fact that it is not directly involved in DNA recognition and binding.^44^ In addition, we also modified *in silico* the DNA lesion to include C<>T, T<>C, and C<>C, as illustrated in Figure 1D, and we also prepared simulation boxes of the same damaged oligonucleotides without the DDB2 unit. To preserve Watson-Crick coupling the base facing the CPD was also modified from A to G when exchanging Thymine with Cytosine. Our MD simulations, which have been performed in replicates to assure a significant statistical sampling, unravel, at an atomistic level, the reasons underscoring the differential repair rates observed experimentally, showing the potentiality of molecular modelling and simulation in addressing relevant (photo-)biological questions.

## COMPUTATIONAL METHODOLOGY

MD simulations were performed on a DNA–protein system derived from the crystallographic structure deposited in the Protein Data Bank (PDB code 4A09),^30^ obtained by X-ray diffraction at a resolution of 3.10 Å. The structure contains a 15-bp oligonucleotide harboring a cyclobutane pyrimidine dimer (CPD) T<>T lesion in complex with the DDB1–DDB2 repair complex. In this study, the DDB1 subunit was removed to focus exclusively on the interactions between DDB2 and the damaged DNA, as DDB1 is not directly involved in lesion recognition and exhibits higher conformational flexibility.

Four DNA systems containing T<>T, T<>C, C<>T, and C<>C CPD lesions were constructed, starting from the crystallographic template. The systems were prepared in silico using the tleap module of the AMBER package. ^50,51^ The T<>C, C<>T, and C<>C were generated from the native T<>T system by mutating the damaged bases while preserving the cyclobutane scaffold of the lesion. The bases complementary to the lesion site have also been rebuilt if needed to obtain complete and structurally consistent double-stranded DNA models. Namely a guanine was inserted whenever a cytosine was present in the CPD.

All systems were solvated in truncated octahedral water box. The TIP3P^51^ explicit solvent model was employed, and potassium (K⁺) counterions were added to ensure the overall electroneutrality of each simulation box.

The DNA/protein systems were parameterized using a combination of force fields. Protein residues were described using the ff14SB force field,^52^ while DNA was modeled using parmBSC1.^53^ Non-standard components, i.e. the CPD lesions, were obtained using the standard Amber procedure and were based on the Amber force Field. Namely the residue point-charges have been obtained using the restraining electrostatic potential (RESP)^54^ by fitting the electrostatic potential obtained from a Harthree-Fock calculation with the 6-31G* basis set (the set of charges have been provided in SI). This procedure has been repeated for each lesion type (T<>T, T<>C, C<>T, and C<>C), ensuring a consistent but lesion-specific force field description across all systems. Note that a specific residue was prepared for the 3’- and 5’ residues. Covalent bonds between the C5–C5 and C6–C6 atoms of the damaged pyrimidines have been enforced using the tleap bonding commands.^50,55^

In addition, four system including DNA only have also been build from the experimental crystal structure following the same protocol and removing entirely the protein.

Hydrogen Mass Repartitioning (HMR)^56^ was consistently applied to enable the use of an increased integration time step of 4 fs. In addition, all bonds involving hydrogen atoms were constrained using the SHAKE algorithm^57^ during molecular dynamics simulations, ensuring numerical stability of the trajectories.

MD simulations were then performed using Amber 24^55^ and a standard multi-step protocol. An initial energy minimization was carried out in two stages. First, positional restraints were applied on the protein while allowing relaxation of solvent and ions, using up to 10,000 cycles including 5,000 steepest descent cycles, with a 10.0 Å non-bonded cutoff. Afterwards, the entire system was fully minimized without restraints using 20,000 cycles, including 10,000 steepest descent cycles (ncyc = 10000), with the same cutoff. The systems were then gradually heated from 0 to 298 K for 2 ns (500,000 steps) under constant volume conditions (NVT ensemble), and without positional constraints. The temperature control was achieved using a Langevin thermostat.^58^ Subsequently, systems were equilibrated during 1 ns (250,000 steps) at 298 K under constant pressure and temperature conditions (NPT ensemble). Pressure conservation was enforced using a Langevin Barostat.^59^ Finally, production MD simulations were carried out under NPT conditions at 298 K and 1 bar using the same thermostat and barostat settings. For each system 4 independent replica of 1 μs were obtained for each system to assure extended statistical sampling.

The trajectory has been visualized using VMD^60^ and analyzed using cpptraj,^61^ also to retrieve the DNA structural parameters with its NucleicAcid plugin, based on the 3DNA approach.^62^ In particular, the opening parameter is defined the signed angular displacement between the x-axes of two DNA base, measured around the average normal to the base-pair plane. The groove parameters have been calculated as the distance between P-P atoms (major) and O4-O4 atoms (minor) as originally proposed by Hossan and Caladine,^63^

## RESULTS

The formation of the complex between DNA and DDB2/DDB1 is recognized as the first step of the NER mechanisms acting in the repair of photolesions and most notably base dimers. It may, thus, be regarded as fundamental to discriminate between the different repair rates observed in skin cells. The action of the DDB2/DDB1 is exerted by its ability to flip the extruded nucleotides out of the double helical arrangements, as also identified by the experimental available crystal structure^30^ visualized in Figure 1C. Importantly the nucleotide excision also induces a more global deformation of the DNA strand, resulting in a visible bump, which facilitates the recruitment of the other NER enzymes, notably the XPC-RAD23B complex, and hence triggers the repair cascade. While this process is relatively well characterized for the T<>T lesion, no experimental structure is available for the lesions involving cytosine. To analyze the effects due to the presence of CPDs, we first report the behavior of solvated DNA strands containing all the four lesions. Note that we have decided to start by the structure of the DNA present in the resolved crystal complex removing the protein and eventually performing mutations to insert C, and G as well as the complementary base, therefore the initial structure of the MD simulation presented extruded dimers. However, upon the very first tens of nanoseconds, the DNA lesions spontaneously insert back in an almost ideal Watson-Crick coupling, which then persists up to the end of the MD simulation. Importantly this behavior is coherent among all the replicas.

This fact can be evidenced pictorially in Figure 2 where we report representative snapshots extracted from the MD simulation in which we can appreciate the perfect insertion of the nucleotide in the Watson and Crick coupling. From a more quantitative point of view, the insertion of the damaged nucleotide can be evidenced by the Opening parameter for the damaged nucleobases. As shown in Figure S3 and S4 we may see that for all the DNA strand the distribution of the opening for the ensemble of the replicas presents an ideal Gaussian shape for all the damaged DNA strands. In all the cases the distribution maximum is found at around 0°, which is indicative of the lesioned dimer being inserted into the strand. Interestingly, all the helical parameters are almost ideal for the four double strands, and particularly the base opening angle parameters describing the extrusion of the nucleotides.

**Figure 2.**
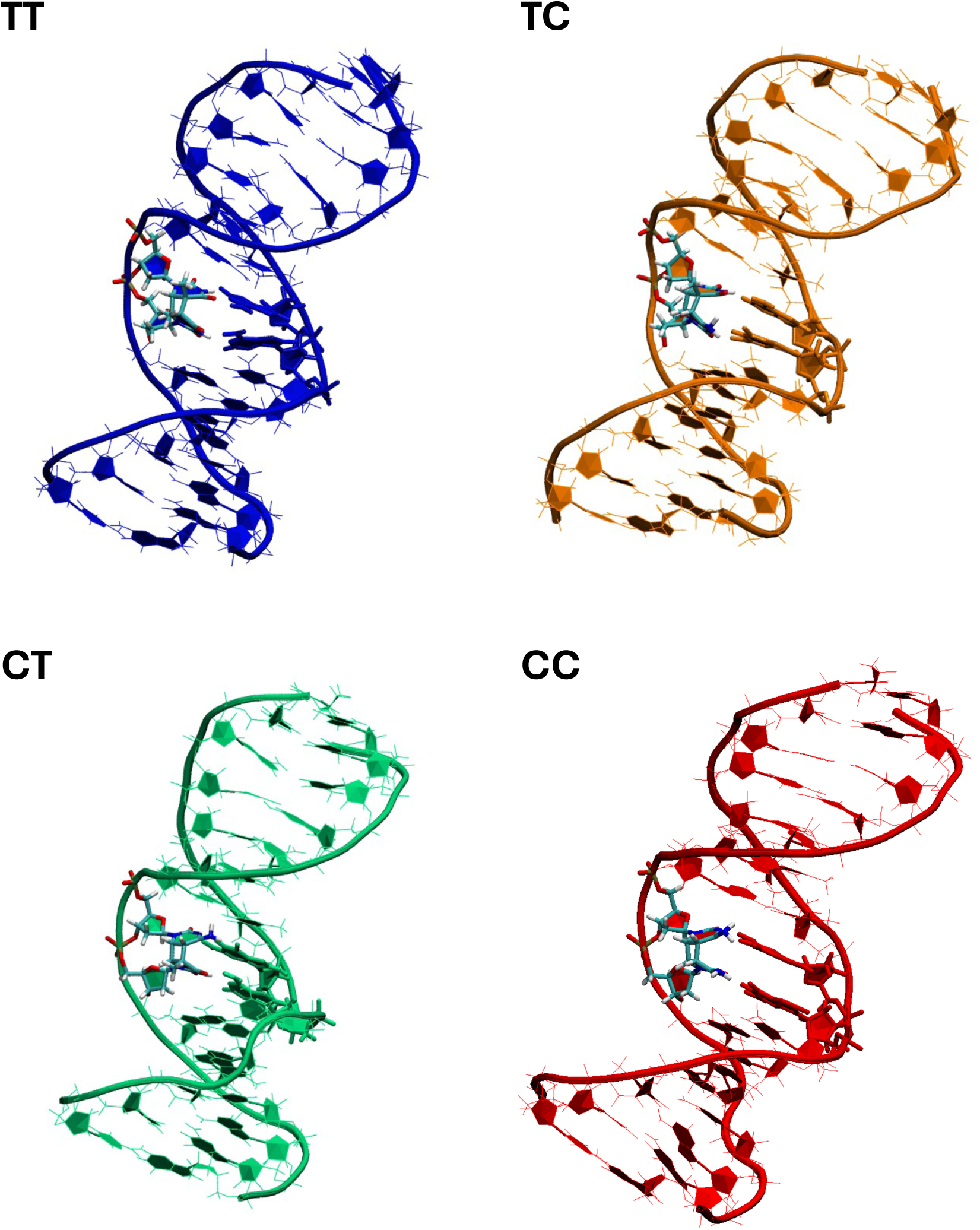
Representative snapshot extracted from the MD simulation for a solvated DNA strand containing the T<>T (blue), T<>C (orange), C<>T (green), and C<>C (blue) lesion, respectively. Note that the same color code will be used throughout the article. The damaged nucleotides are highlighted in licorice representation.

Globally, the structural modifications observed in the case of the naked DNA strand are not sufficient to highlight significant differences, which may justify the experimentally observed repair rates among the different lesions, and in particular the better repair propensity when a cytosine is present at the 5’ position of the CPD. Therefore, we have performed MD simulations on the DDB2/DNA complex bearing the four CPD lesions for which representative snapshots are reported in Figure 3.

**Figure 3.**
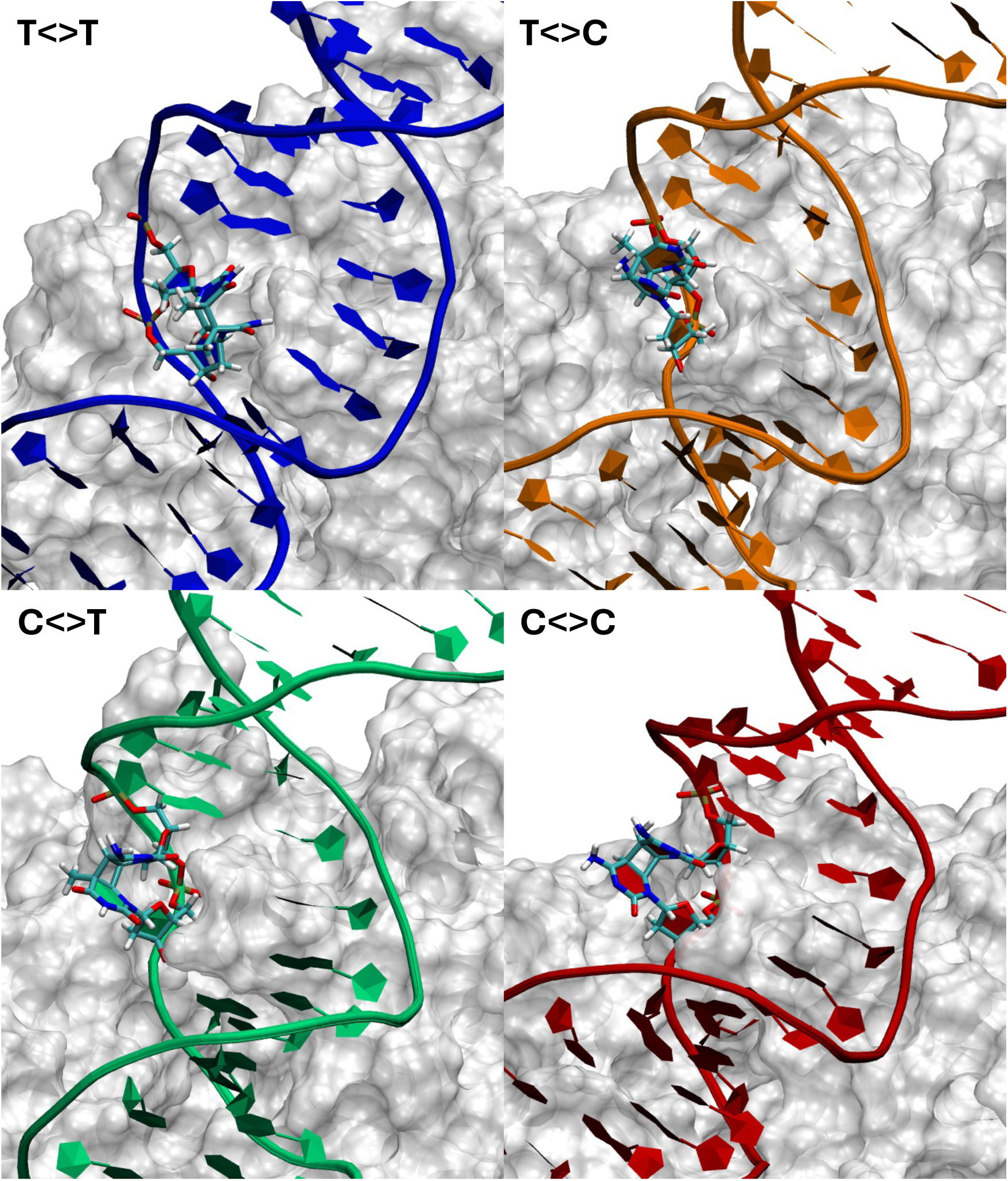
Representative snapshot for the interaction between DDB2 and a DNA strand containing T<>T, T<>C, C<>T, and C<>C, respectively. DNA is represented in cartoon, the CPD in licorice and the protein as a surface.

For all the lesions (Figure 3), the CPD damage is globally extruded coherently with what observed for the crystal structure of the T<>T lesion and with the biological role played by DDB2 in favoring the recognition of CPDs. We may also observe that, for all the studied lesions, the space left by the extruded CPD in the DNA helix is compensated by a partial intercalation of a triad of hydrophobic or polar aminoacid, namely, H273, Q272, and F271. The preeminent role of these aminoacids in shaping the interaction network leading to CPD extrusion will be discussed later.

However, this apparent straightforward structure masks a more complex behavior and some subtle differences between the strands. Indeed, if we consider the distribution of the base Opening angle between the damaged bases and the complementary ones (Figure 4 and S5), we may see that while T<>T and C<>C show stable extruded structure over the ensemble of the MD simulation with average values centered around 100°, T<>C and C<>T present a bimodal distribution in which the extruded structure is accompanied by a conformation characterized by much smaller Opening centered at about 10° (C<>T) and 30° (T<>C), respectively. Clearly, this corresponds to conformations in which the extrusion of the damage nucleobase is partially lost. As shown in Figure 5, they are characterized by a rotation of the backbone dihedral which reorients the damaged dimers towards the DNA core. Furthermore, the interaction with the H273, Q272, and F271 triad is also reshaped. More importantly, while the damaged nucleobases are clearly reoriented towards the DNA core, the ideal Watson-Crick pairing is not reinstated and the DNA structure appears mismatched, maintaining a considerable structural deformation which may still be compatible with the recognition by the downstream repair complexes.

**Figure 4.**
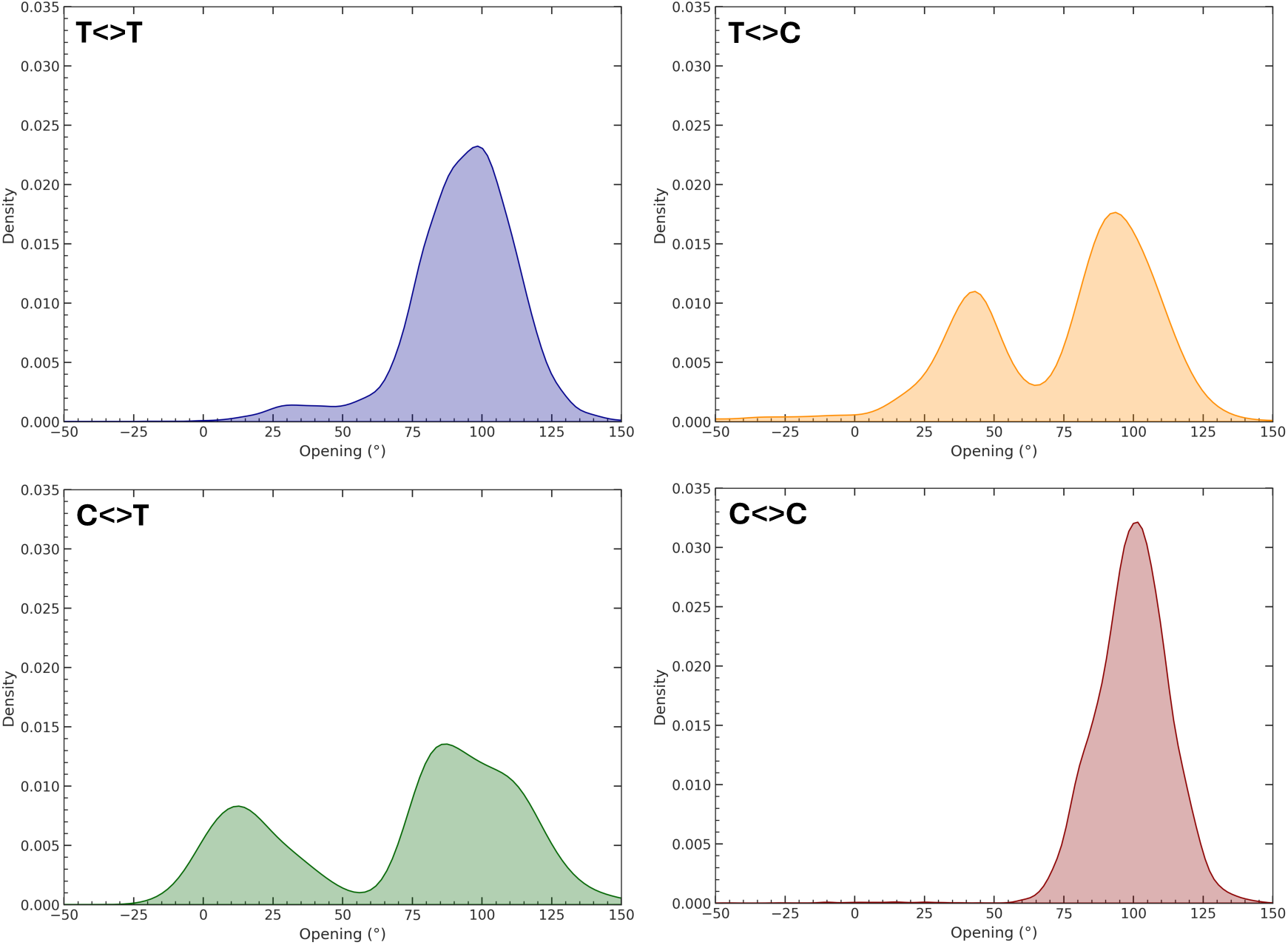
Distribution of Opening for the 5’ damaged nucleotide on the ensemble of the DNA/DDB2 trajectories. The corresponding distribution for the 3’ nucleotide is provided in the SI (Figure S5) and follows the exact tendencies.

**Figure 5.**
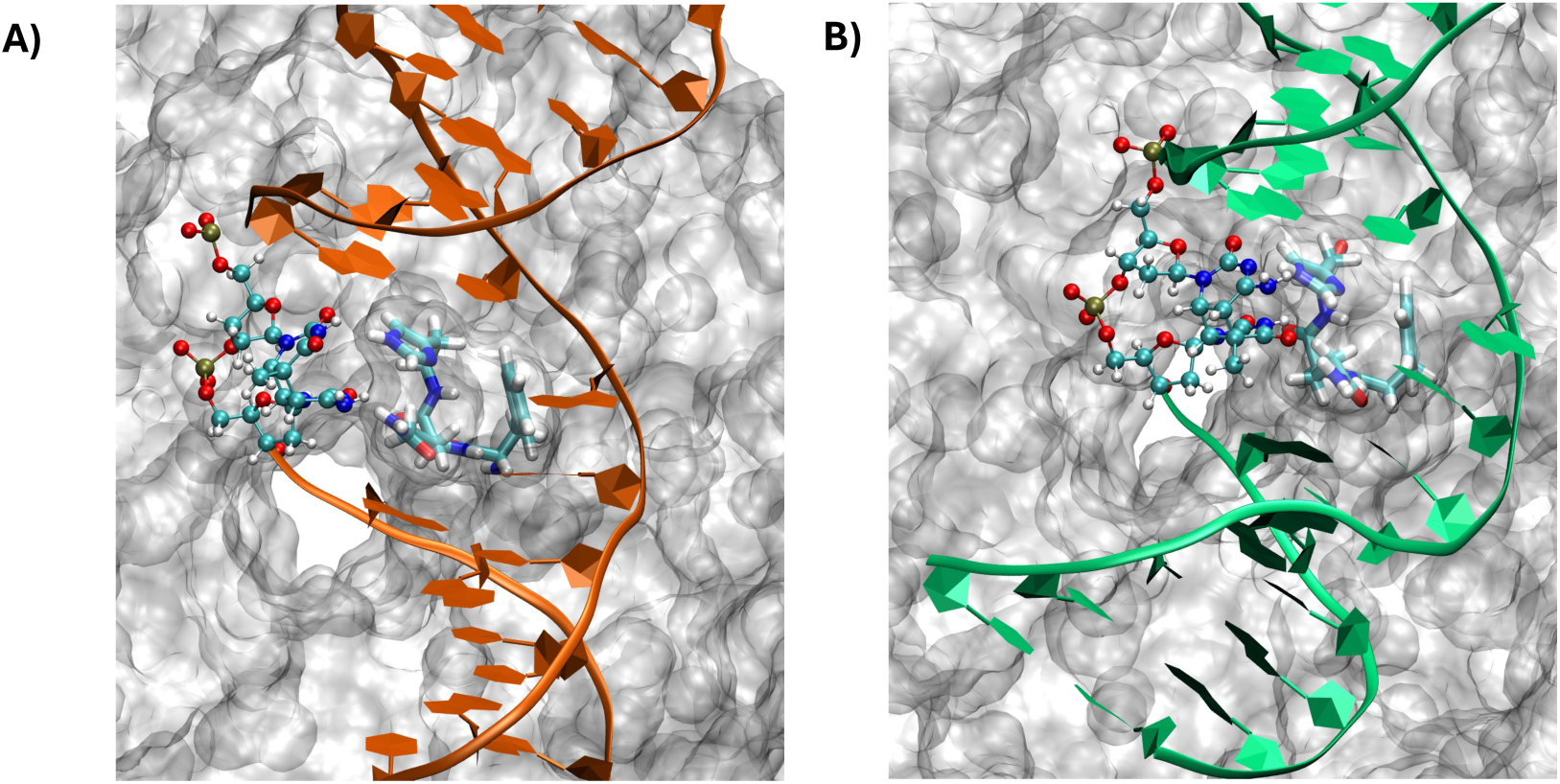
Representative snapshot showing the non-extruded conformation observed for T<>C (A) and C<>T (B). The damaged nucleotides are represented in ball and stick, while the H273, Q272, and F271triad is shown in licorice representation.

Despite this important difference in the behavior of the damaged strands and considering the presence of either T<>T and C<>C or instead T<>C and C<>T lesions, the correlation between the structural parameters and CPD repair rate is still lacking. However, we should recall that the complex NER pathway involves the recruitment of the XPC-RAD23B complex downstream of the binding with DDB2. Indeed, XPC can recognize structurally deformed DNA strands bearing the landmarks of the presence of lesions. The action of DDB2 extruding the damaged nucleobases and deforming the DNA is, thus, instrumental in allowing further recognition. As a matter of fact, XPC appears to sense the DNA groove parameters and particularly the complementary strand which is not directly interacting with DDB2 and may be accessed more easily. Since XPC mainly recognizes the deformations of the grooves taking place in the immediate proximity of the CPD lesion we have assessed the distribution of the DNA major groove width at the -1 and +1 positions with respect to the CPD (Figure 6 and Figure S6). The deformations of the minor groove, which are reported in SI only (Figures S7 and S8), are less relevant since the latter is engaged in the interaction with DDB2 and cannot be sensed by the downstream repair complexes.

**Figure 6.**
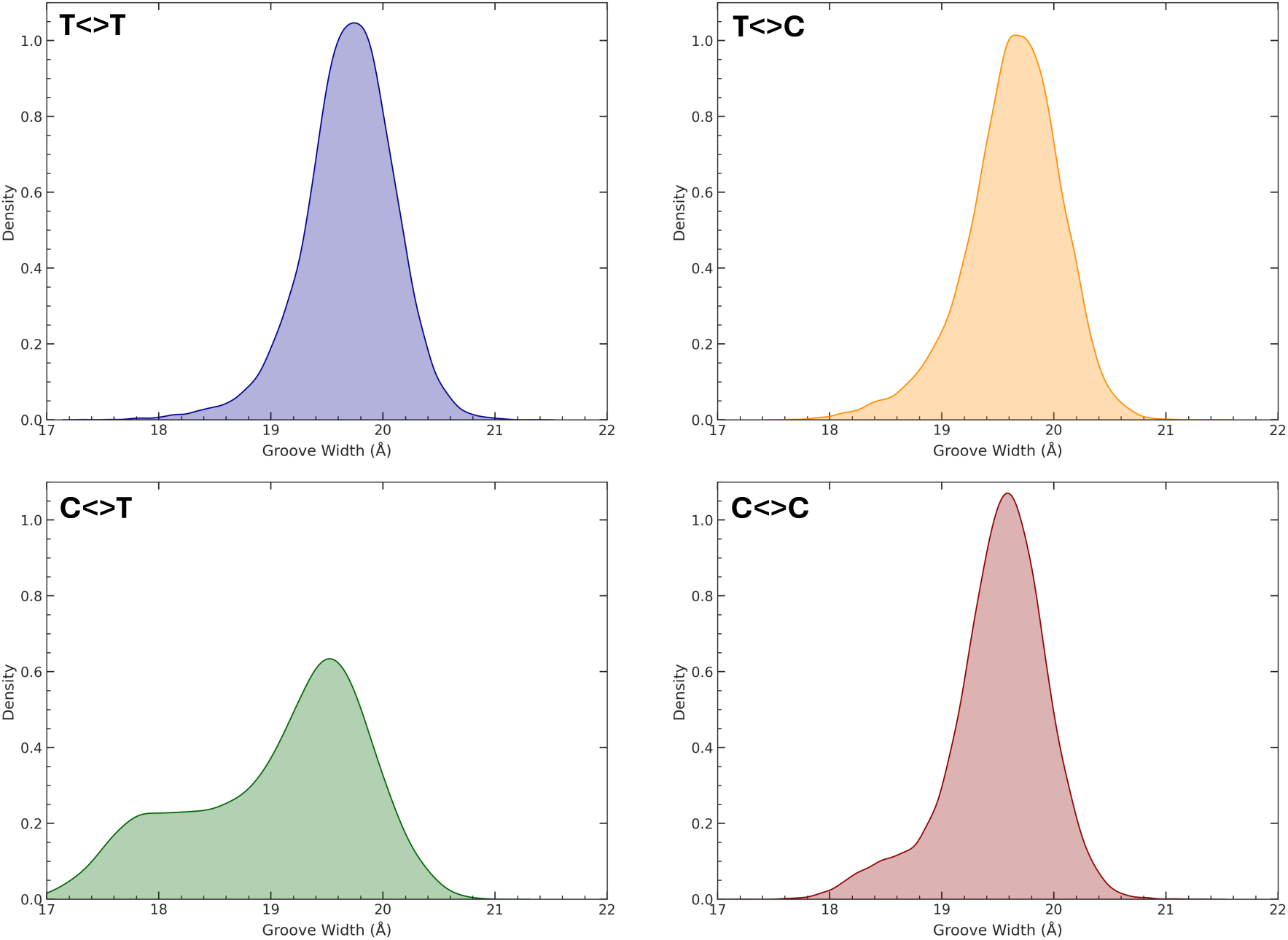
Distribution of the major groove width as extracted from the MD trajectory at the position -1 with respect to the CPD lesion, the corresponding width for the +1 position are reported in SI.

In ideal B-DNA the major groove width should peak around 21 Å, with some slight variations depending on the specific sequence. We may see that upon the binding with DDB2 and for both the -1 (Figure 6) and +1 (Figure S6) positions the major groove is instead slightly shrunk, which is a clear signature of a structural deformation showing a principal maximum at about 19.5 Å. While no significant differences among the strands may be observed for the +1 position interesting features are instead emerging at the -1 position. Indeed, T<>T and T<>C presents a well-defined, almost perfect Gaussian distribution, while larger tail appears for C<>T and at least partially C<>C. In the latter case, an evident shoulder in the distribution is present, which implies the population of groove widths reaching about 18 Å. While the shoulder is only scarcely evident in the case of C<>C, C<>T presents an extended plateau at the small width regions. Thus, while the extrusion of the DNA lesion is almost equivalent whatever the base forming the CPD damage, the presence of a cytosine in 5’ position is favoring a more pronounced deformation of the complementary strand which is reflected in the groove structural parameters. The larger deformation of the DNA, compared to the canonical B-DNA form may then favor the interaction with the XPC repair protein, which should sense large strand deformation, and, consequently, may lead to a more efficient repair. The significant difference in the distribution of the -1 major width can be better appreciated by using a Gaussian Mixture Model analysis. Indeed, for all the lesions the distribution may be represented by two gaussian functions as shown in Figure S13 and Table S1. The largest weight components is always cantered at around 19.5-19.8 Å, while the second component almost totally overlaps with the first one in the case of T<>T and is instead centred at 18.1 Å for the T<>C lesion. For T<>C and C<>C a slightly intermediate situation holds. However, a drift of the second component mean value can be observed for C<>C coherently with the repair rate.

The amount of the deformation of the complementary strand and the major groove may reflect the repair rate efficiency, with the C<>T damage being the most efficiently repair and presenting the most important deformation. Interestingly, the deformation of the major groove parameters at the lesion site are less significant both in terms of their difference with respect to the canonical B-DNA and between the damages (Figures S9-S12).

To better understand the effects indued by the binding to DDB2 we also report the most crucial interaction at a residue level between the protein aminoacids and the nucleic acid nucleobases. As already described, the DDB2 triad comprised of H273, Q272, and F271 are the most important residues stabilizing the extruded base, and this whatever the nature of the pyrimidine engaged in the DNA damage.

At the residue level, we may observe that a complex interaction network between the protein and the damaged DNA strand is in place. Not surprisingly, the DNA backbone is stabilized by a series of basic residues leading to an accumulation of positive charges interacting favorably with the deprotonated phosphate groups. At the CPD site, coherently with what is reported in the crystal structure, we may identify R48, R49, and K68 as the most important residues. In Figure 7 we report the distribution of the distances of these residues with the closest phosphate groups along the MD simulations for the four systems, confirming the persistence of the main interactions, and especially those involving R48 and K68, for T<>T and C<>C. While the interaction pattern is globally confirmed in the case of T<>C and C<>T strands, we may evidence partial losses of the interactions particularly involving R49 and to a less extent K68.

**Figure 7.**
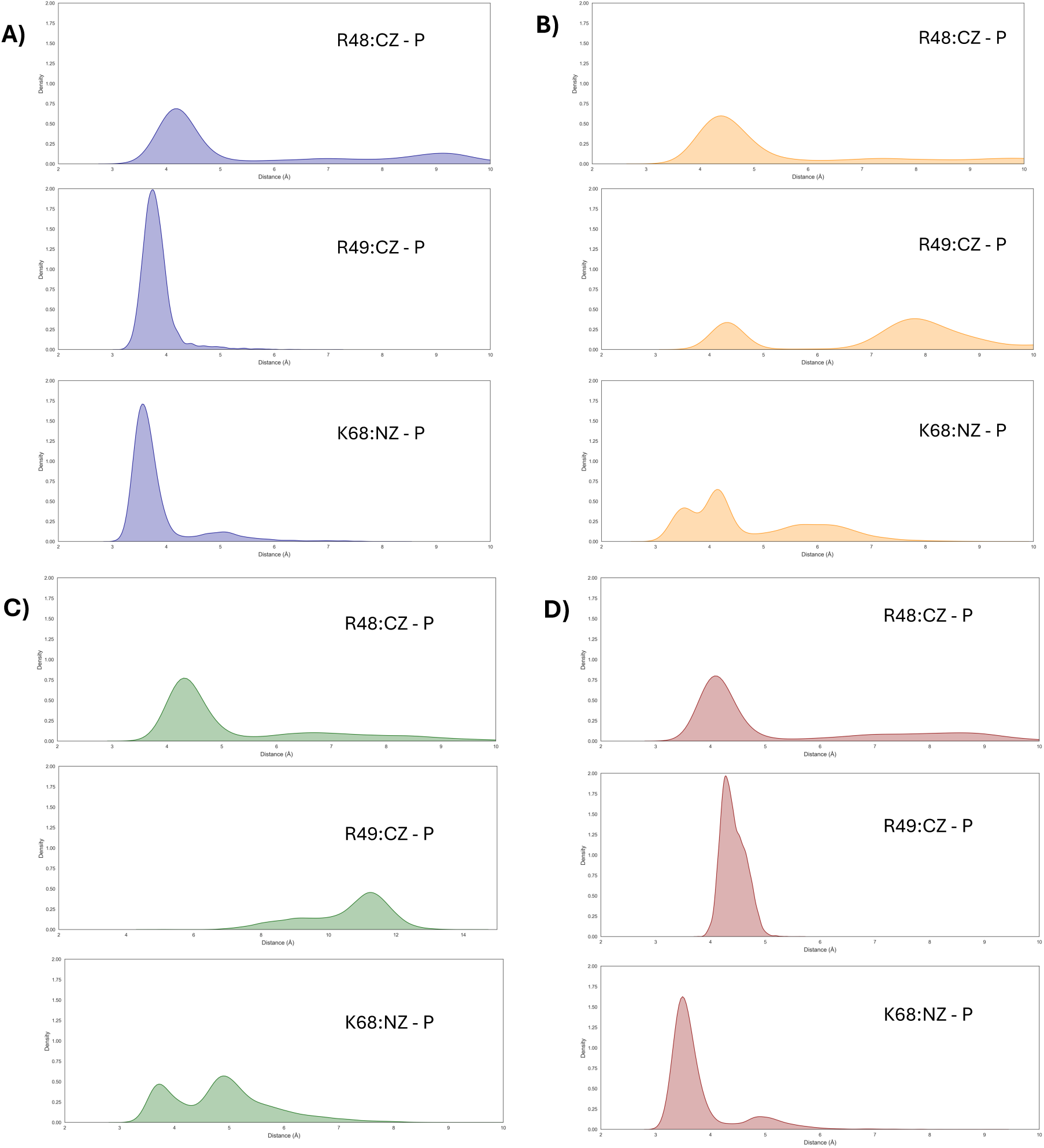
Distribution of the distance between the DNA backbone phosphate and R48, R49, and K68 for T<>T (A), T<>C (B), C<>T (C), and C<>C (D), respectively. The distances have been calculated considering the CZ atom of arginine and NZ in the case of lysine with respect to the backbone P atom.

Interestingly the analysis of the time series confirms that the reshaping of the interaction network correlates with the reinsertion of the damaged dinucleotide in the DNA helical structure, which has been observed for these two systems. In the case of C<>T the interaction involving R49 is particularly weakened leading to very large distance distribution extending up to 12 Å. As shown in SI, when the salt bridge with the DNA backbone is broken R49 develops a hydrogen bond with the backbone of the H273 leading to a stable and rather persistent secondary conformation.

As previously described, H273 and Q272 have been pointed out as fundamental to stabilize the extruded CPD lesion and, at least partially, compensate for the loss of Watson-Crick hydrogen bonds. Therefore, in Figure 8 we report the distribution of the distances of the two residues with the pyrimidine carbonyl moiety. Interestingly, H273 develops preferential interactions with the 5’ nucleobases, while Q272 is mainly locking the 3’ residue.

**Figure 8.**
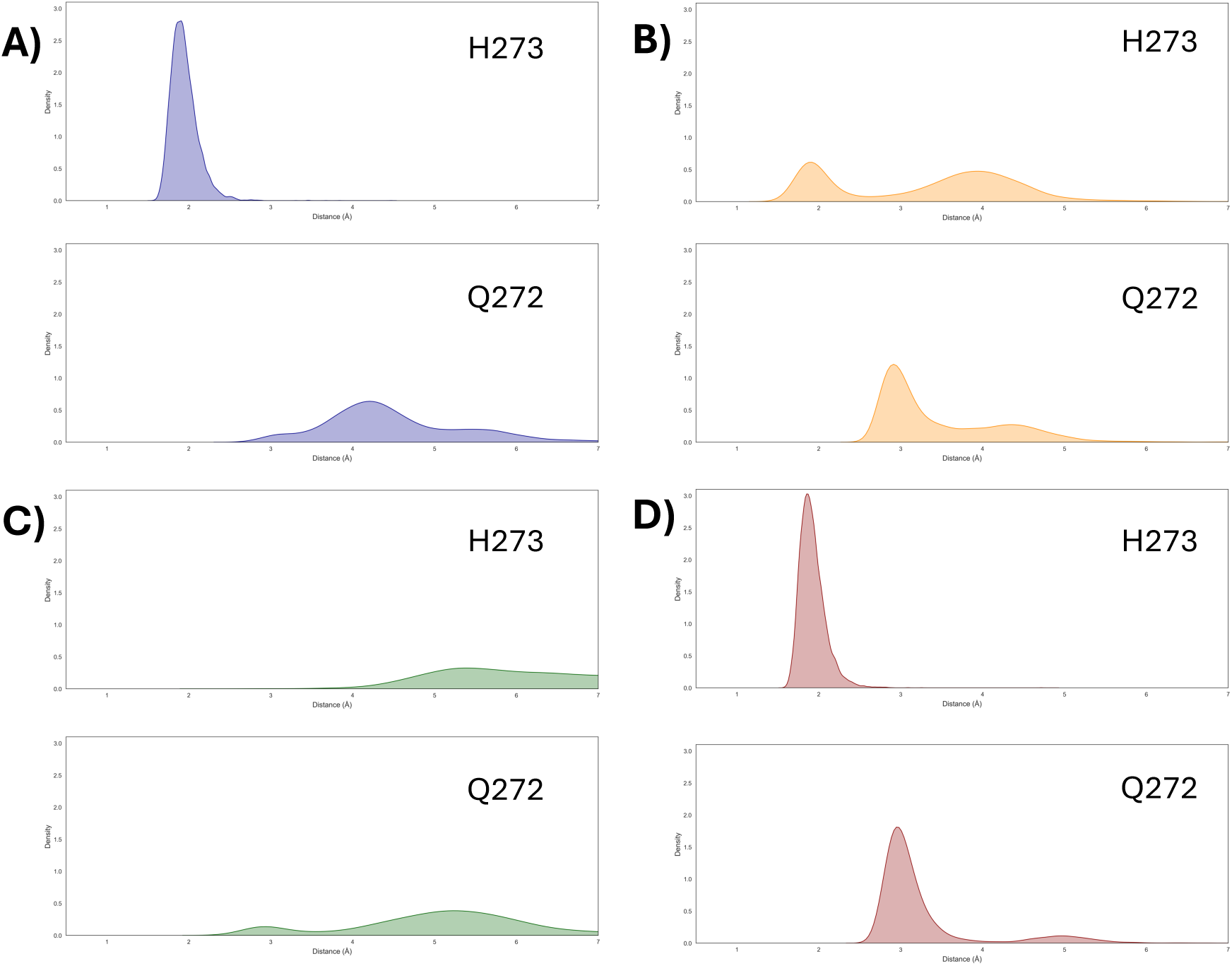
Distribution of the distance between the lesioned nucleotides and H273 and Q272 for T<>T (A), T<>C (B), C<>T (C), and C<>C (D), respectively.

A crucial difference may be observed in the four damaged systems. Indeed, while the hydrogen bond is extremely stable and persistent in both T<>T and C<>C strand it is again partially lost in the case of the heterodimers. This is also particularly pronounced for the C<>T strand and may be ascribed to the formation of the secondary hydrogen bond between H273 and R48 as previously discussed. However, the observed differences, while they cannot be directly related to the repair rate, point again to the crucial role of these two residues in favoring conformations with extruded nucleobases. Indeed, the loss of the interaction with H273 and Q272 correlates well with the reinsertion of the damaged CPD.

## CONCLUSIONS

CPD lesions accumulate into DNA due to their general bad repair. Yet, its efficiency depends on the nature of the damage, the presence of a cytosine in 5’ position being beneficial for triggering the activity of the NER pathway. By performing all-atom MD simulations, we have sketched a plausible correlation between the differential repair rate of CPD DNA lesions, and the structural deformations induced upon the interaction with DDB2, which represents the first event of the NER pathway. We have shown that no significant structural deformation of the DNA lesions or deviation from the ideal canonical structure may be observed, whatever the specific CPD formed. This observation, which is coherent with a large literature, also justifies the difficulty of the repair of CPD lesions which tend to accumulate in cells. The interaction with the DDB1/DDB2 repair complex is, instead, inducing strong modification of the helical structure, which most notably the extrusion of the CPD damage, as also confirmed by crystal structure for the T<>T containing strand.^30^ However, no significant difference on the interactions developed with the DDB2 protein or on the damaged basis extrusion. Conversely, we have shown that C<>C and, more significantly C<>T, lesions are presenting a much larger structural deformation of the complementary strand compared to T<>C and C<>T, as particularly reflected on the major groove width parameters. In turn, the more pronounced structural deformation may be related to a better recognition by the downstream XPC-RAD23B complex, which should bind to the damaged DNA to finalize the repair. While our results do not allow for a quantitative correlation and reproduction of the different repair rates, they tend to draw important tendencies which point to a different structural deformation as a function of the specific type of CPD, also showing some of the capabilities of *in silico* photobiology.

However, in the future, we plan to explicitly explore the binding of XPC to the DDB2/DNA complex, eventually estimating the complexation free energy, to precisely assess the role of the deformation of the complementary strand in promoting DNA repair.

## ACKNOWLEDGEMENTS

The authors thank GENCI and Explor, for computational resources. The authors thanks ANR and CGI for their financial support of this work through Labex SEAM ANR 11 LABEX 086. The support of the IdEx “Université Paris 2019” ANR-18-IDEX-0001. Support from the PEPR LUMA MANDALA is also gratefully acknowledged.

## REFERENCES

(1) Ikehata, H.; Ono, T. The Mechanisms of UV Mutagenesis. J. Radiat. Res. 2011, 52 (2), 115–125. 10.1269/jrr.10175.

(2) Rünger, T. M.; Kappes, U. P. Mechanisms of Mutation Formation with Long-Wave Ultraviolet Light (UVA). Photodermatology Photoimmunology and Photomedicine. Blackwell Publishing Ltd 2008, pp 2–10. 10.1111/j.1600-0781.2008.00319.x.

(3) Cadet, J.; Mouret, S.; Ravanat, J. L.; Douki, T. Photoinduced Damage to Cellular DNA: Direct and Photosensitized Reactions. In Photochemistry and Photobiology; Blackwell Publishing Ltd, 2012; Vol. 88, pp 1048–1065. 10.1111/j.1751-1097.2012.01200.x.

(4) Cadet, J.; Douki, T.; Ravanat, J. L. Oxidatively Generated Damage to Cellular DNA by UVB and UVA Radiation. Photochemistry and Photobiology. 2015, pp 140–155. 10.1111/php.12368.

(5) Drouin, R.; Therrien, J. P. UVB-Induced Cyclobutane Pyrimidine Dimer Frequency Correlates with Skin Cancer Mutational Hotspots in P53. Photochem. Photobiol. 1997, 66 (5), 719–726. 10.1111/j.1751-1097.1997.tb03213.x.

(6) Sage, E.; Lamolet, B.; Brulay, E.; Moustacchi, E.; Chteauneuf, A.; Drobetsky, E. A. Mutagenic Specificity of Solar UV Light in Nucleotide Excision Repair-Deficient Rodent Cells. Proc. Natl. Acad. Sci. U. S. A. 1996, 93 (1), 176–180. 10.1073/pnas.93.1.176.

(7) Pfeifer, G. P.; You, Y. H.; Besaratinia, A. Mutations Induced by Ultraviolet Light. Mutation Research - Fundamental and Molecular Mechanisms of Mutagenesis. Elsevier April 1, 2005, pp 19–31. 10.1016/j.mrfmmm.2004.06.057.

(8) Sage, E. Distribution and Repair of Photolesions in DNA: Genetic Consequences and the Role of Sequence Context. Photochem. Photobiol. 1993, 57 (1), 163–174. 10.1111/j.1751-1097.1993.tb02273.x.

(9) Gustavsson, T.; Improta, R.; Markovitsi, D. DNA/RNA: Building Blocks of Life under UV Irradiation. Journal of Physical Chemistry Letters. 2010, pp 2025–2030. 10.1021/jz1004973.

(10) Bhagavan, N. V.; Ha, C.-E. Structure and Properties of DNA Photoproducts. In Essentials of Medical Biochemistry; CRC Press, 2015; pp 381–400. 10.1016/B978-0-12-416687-5.00021-X.

(11) Cadet, J.; Sage, E.; Douki, T. Ultraviolet Radiation-Mediated Damage to Cellular DNA. Mut. Res. Fund. Mol. Mec. Mut. 2005, 571 (1–2), 3–17. 10.1016/j.mrfmmm.2004.09.012.

(12) Banyasz, A.; Douki, T.; Improta, R.; Gustavsson, T.; Onidas, D.; Vayá, I.; Perron, M.; Markovitsi, D. Electronic Excited States Responsible for Dimer Formation upon UV Absorption Directly by Thymine Strands: Joint Experimental and Theoretical Study. J. Am. Chem. Soc. 2012, 134 (36), 14834–14845. 10.1021/ja304069f.

(13) Rauer, C.; Nogueira, J. J.; Marquetand, P.; González, L. Cyclobutane Thymine Photodimerization Mechanism Revealed by Nonadiabatic Molecular Dynamics. J. Am. Chem. Soc. 2016, 138 (49), 15911–15916. 10.1021/jacs.6b06701.

(14) Epe, B. DNA Damage Spectra Induced by Photosensitization. Photochem. Photobiol. Sci. 2012, 11 (1), 98–106. 10.1039/C1PP05190C.

(15) Cuquerella, M. C.; Lhiaubet-Vallet, V.; Cadet, J.; Miranda, M. A. Benzophenone Photosensitized DNA Damage. Acc. Chem. Res. 2012, 45 (9), 1558–1570. 10.1021/ar300054e.

(16) Dumont, E.; Wibowo, M.; Roca-Sanjuán, D.; Garavelli, M.; Assfeld, X.; Monari, A. Resolving the Benzophenone DNA-Photosensitization Mechanism at QM/MM Level. Journal of Physical Chemistry Letters 2015, 6 (4), 576–580. 10.1021/jz502562d.

(17) Gentile, M.; Talotta, F.; Tremblay, J. C.; González, L.; Monari, A. Predominant Binding Mode of Palmatine to DNA. J. Phys. Chem. Lett. 2024, 15 (42), 10570– 10575. 10.1021/acs.jpclett.4c02721.

(18) Sinha, R. P.; Häder, D.-P. UV-Induced DNA Damage and Repair: A Review. Photochemical & Photobiological Sciences 2002, 1 (4), 225–236. 10.1039/b201230h.

(19) Kim, J. -K; Patel, D.; Choi, B. -S. Contrasting Structural Impacts Induced by Cis-syn Cyclobutane Dimer and (6–4) Adducts in DNA Duplex Decamers: Implication in Mutagenesis and Repair Activity. Photochem. Photobiol. 1995, 62 (1), 44–50. 10.1111/j.1751-1097.1995.tb05236.x.

(20) Kim, J. -K; Choi, B. -S. The Solution Structure of DNA Duplex-Decamer Containing the (6-4) Photoproduct of Thymidylyl(3′→5′)Thymidine by NMR and Relaxation Matrix Refinement. Eur. J. Biochem. 1995, 228 (3), 849–854. 10.1111/j.1432-1033.1995.0849m.x.

(21) Jing, Y.; Kao, J. F.-L.; Taylor, J. S. Thermodynamic and Base-Pairing Studies of Matched and Mismatched DNA Dodecamer Duplexes Containing Cis-Syn, (6-4) and Dewar Photoproducts of TT. Nucleic Acids Res. 1998, 26 (16), 3845–3853. 10.1093/nar/26.16.3845.

(22) Fujiwara, Y.; Iwai, S. Thermodynamic Studies of the Hybridization Properties of Photolesions in DNA. Biochemistry 1997, 36 (6), 1544–1550. 10.1021/bi9619942.

(23) Park, C. J.; Lee, J. H.; Choi, B. S. Functional Insights Gained from Structural Analyses of DNA Duplexes That Contain UV-Damaged Photoproducts. Photochem. Photobiol. 2007, 83 (1), 187–195. 10.1562/2006-02-28-IR-820.

(24) Dehez, F.; Gattuso, H.; Bignon, E.; Morell, C.; Dumont, E.; Monari, A. Conformational Polymorphism or Structural Invariance in DNA Photoinduced Lesions: Implications for Repair Rates. Nucleic Acids Res. 2017, 45 (7), 3654–3662. 10.1093/nar/gkx148.

(25) Sedletska, Y.; Radicella, J. P.; Sage, E. Replication Fork Collapse Is a Major Cause of the High Mutation Frequency at Three-Base Lesion Clusters. Nucleic Acids Res. 2013, 41 (20), 9339–9348. 10.1093/nar/gkt731.

(26) Sastry, M.; Patel, D. J. Solution Structure of the Mithramycin Dimer-DNA Complex. Biochemistry 1993, 32 (26), 6588–6604.

(27) Johnson, R. E.; Haracska, L.; Prakash, S.; Prakash, L. Role of DNA Polymerase Eta in the Bypass of a (6-4) TT Photoproduct. Mol. Cell. Biol. 2001, 21 (10), 3558–3563. 10.1128/MCB.21.10.3558-3563.2001.

(28) Pfeifer, G. P. Formation and Processing of UV Photoproducts: Effects of DNA Sequence and Chromatin Environment. Photochem. Photobiol. 1997, 65 (2), 270–283. 10.1111/j.1751-1097.1997.tb08560.x.

(29) Mouret, S.; Charveron, M.; Favier, A.; Cadet, J.; Douki, T. Differential Repair of UVB-Induced Cyclobutane Pyrimidine Dimers in Cultured Human Skin Cells and Whole Human Skin. DNA Repair (Amst). 2008, 7 (5), 704–712. 10.1016/j.dnarep.2008.01.005.

(30) Scrima, A.; Koníčková, R.; Czyzewski, B. K.; Kawasaki, Y.; Jeffrey, P. D.; Groisman, R.; Nakatani, Y.; Iwai, S.; Pavletich, N. P.; Thomä, N. H. Structural Basis of UV DNA-Damage Recognition by the DDB1-DDB2 Complex. Cell 2008, 135 (7), 1213–1223. 10.1016/j.cell.2008.10.045.

(31) Ziegler, A.; Leffell, D. J.; Kunala, S.; Sharma, H. W.; Gailani, M.; Simon, J. A.; Halperin, A. J.; Baden, H. P.; Shapiro, P. E.; Bale, A. E. Mutation Hotspots Due to Sunlight in the P53 Gene of Nonmelanoma Skin Cancers. Proc. Natl. Acad. Sci. U. S. A. 1993, 90 (9), 4216–4220. 10.1073/pnas.90.9.4216.

(32) Brash, D. E.; Rudolph, J. A.; Simon, J. A.; Lin, A.; McKenna, G. J.; Baden, H. P.; Halperin, A. J.; Pontén, J. A Role for Sunlight in Skin Cancer: UV-Induced P53 Mutations in Squamous Cell Carcinoma. Proc. Natl. Acad. Sci. U. S. A. 1991, 88 (22), 10124–10128.

(33) Lawrence, C. W.; Gibbs, P. E. M.; Borden, A.; Horsfall, M. J.; Kilbey, B. J. Mutagenesis Induced by Single UV Photoproducts in E. Coli and Yeast. Mutation Research/Genetic Toxicology 1993, 299 (3–4), 157–163. 10.1016/0165-1218(93)90093-S.

(34) Taylor, J. S. Unraveling the Molecular Pathway from Sunlight to Skin Cancer. Acc. Chem. Res. 1994, 27 (3), 76–82. 10.1021/ar00039a003.

(35) Cole, C.; Forbes, P. D. Examining the Role of Visible Light in Photocarcinogenesis – Lessons from the Past. *J*. Photochem. Photobiol. 2023, 17, 100201. 10.1016/j.jpap.2023.100201.

(36) Perilla, J. R.; Goh, B. C.; Cassidy, C. K.; Liu, B.; Bernardi, R. C.; Rudack, T.; Yu, H.; Wu, Z.; Schulten, K. Molecular Dynamics Simulations of Large Macromolecular Complexes. Current Opinion in Structural Biology. April 2015, pp 64–74. 10.1016/j.sbi.2015.03.007.

(37) Dumont, E.; Monari, A. Understanding DNA under Oxidative Stress and Sensitization: The Role of Molecular Modeling. Front. Chem. 2015, 3, 43. 10.3389/fchem.2015.00043.

(38) Gattuso, H.; Durand, E.; Bignon, E.; Morell, C.; Georgakilas, A. G.; Dumont, E.; Chipot, C.; Dehez, F.; Monari, A. Repair Rate of Clustered Abasic DNA Lesions by Human Endonuclease: Molecular Bases of Sequence Specificity. Journal of Physical Chemistry Letters 2016, 7 (19), 3760–3765. 10.1021/acs.jpclett.6b01692.

(39) Karami, Y.; Bignon, E. Cysteine Hyperoxidation Rewires Communication Pathways in the Nucleosome and Destabilizes the Dyad. Comput. Struct. Biotechnol. J. 2024, 23, 1387–1396. 10.1016/j.csbj.2024.03.025.

(40) Matoušková, E.; Bignon, E.; Claerbout, V. E. P.; Dršata, T.; Gillet, N.; Monari, A.; Dumont, E.; Lankaš, F. Impact of the Nucleosome Histone Core on the Structure and Dynamics of DNA-Containing Pyrimidine-Pyrimidone (6-4) Photoproduct. J. Chem. Theory Comput. 2020, 16 (9), 5972–5981. 10.1021/acs.jctc.0c00593.

(41) Francés-Monerris, A.; Gillet, N.; Dumont, E.; Monari, A. DNA Photodamage and Repair: Computational Photobiology in Action. In Challenges and Advances in Computational Chemistry and Physics; Springer Science and Business Media B.V., 2021; Vol. 31, pp 293–332. 10.1007/978-3-030-57721-6_7.

(42) Schärer, O. D. Nucleotide Excision Repair in Eukaryotes. Cold Spring Harbor Perspectives in Biology. Cold Spring Harbor Laboratory Press October 1, 2013, p a012609. 10.1101/cshperspect.a012609.

(43) Dexheimer, T. S. DNA Repair Pathways and Mechanisms. In DNA Repair of Cancer Stem Cells; 2013; pp 19–32. 10.1007/978-94-007-4590-2_2.

(44) Stoyanova, T.; Roy, N.; Kopanja, D.; Raychaudhuri, P.; Bagchi, S. DDB2 (Damaged-DNA Binding Protein 2) in Nucleotide Excision Repair and DNA Damage Response. Cell Cycle 2009, 8 (24), 4067–4071. 10.1016/j.micinf.2011.07.011.

(45) Matsumoto, S.; Fischer, E. S.; Yasuda, T.; Dohmae, N.; Iwai, S.; Mori, T.; Nishi, R.; Yoshino, K. I.; Sakai, W.; Hanaoka, F.; Thomä, N. H.; Sugasawa, K. Functional Regulation of the DNA Damage-Recognition Factor DDB2 by Ubiquitination and Interaction with Xeroderma Pigmentosum Group C Protein. Nucleic Acids Res. 2015, 43 (3), 1700–1713. 10.1093/nar/gkv038.

(46) Tsuge, M.; Masuda, Y.; Kaneoka, H.; Kidani, S.; Miyake, K.; Iijima, S. SUMOylation of Damaged DNA-Binding Protein DDB2. Biochem. Biophys. Res. Commun. 2013, 438 (1), 26–31. 10.1016/j.bbrc.2013.07.013.

(47) Li, J.; Wang, Q. E.; Zhu, Q.; El-Mahdy, M. A.; Wani, G.; Prætorius-Ibba, M.; Wani, A. A. DNA Damage Binding Protein Component DDB1 Participates in Nucleotide Excision Repair through DDB2 DNA-Binding and Cullin 4a Ubiquitin Ligase Activity. Cancer Res. 2006, 66 (17), 8590–8597. 10.1158/0008-5472.CAN-06-1115.

(48) Pines, A.; Vrouwe, M. G.; Marteijn, J. A.; Typas, D.; Luijsterburg, M. S.; Cansoy, M.; Hensbergen, P.; Deelder, A.; de Groot, A.; Matsumoto, S.; Sugasawa, K.; Thoma, N.; Vermeulen, W.; Vrieling, H.; Mullenders, L. PARP1 Promotes Nucleotide Excision Repair through DDB2 Stabilization and Recruitment of ALC1. Journal of Cell Biology 2012, 199 (2), 235–249. 10.1083/jcb.201112132.

(49) An, S.; Kusakabe, M.; Kim, H.-S.; Kozono, H.; Cheon, N. Y.; Kim, J.; Kang, J.; Jang, S.; Sugasawa, K.; Schärer, O. D.; Lee, J. Y. XPC-RAD23B Enhances UV-DDB Binding to DNA to Facilitate Lesion Search in Nucleotide Excision Repair. Nucleic Acids Res. 2025, 53 (11). 10.1093/nar/gkaf463.

(50) Case, D. A.; Aktulga, H. M.; Belfon, K.; Cerutti, D. S.; Cisneros, G. A.; Cruzeiro, V. W. D.; Forouzesh, N.; Giese, T. J.; Götz, A. W.; Gohlke, H.; Izadi, S.; Kasavajhala, K.; Kaymak, M. C.; King, E.; Kurtzman, T.; Lee, T.-S.; Li, P.; Liu, J.; Luchko, T.; Luo, R.; Manathunga, M.; Machado, M. R.; Nguyen, H. M.; O’Hearn, K. A.; Onufriev, A. V.; Pan, F.; Pantano, S.; Qi, R.; Rahnamoun, A.; Risheh, A.; Schott-Verdugo, S.; Shajan, A.; Swails, J.; Wang, J.; Wei, H.; Wu, X.; Wu, Y.; Zhang, S.; Zhao, S.; Zhu, Q.; Cheatham, T. E.; Roe, D. R.; Roitberg, A.; Simmerling, C.; York, D. M.; Nagan, M. C.; Merz, K. M. AmberTools. J. Chem. Inf. Model. 2023, 63 (20), 6183–6191. 10.1021/acs.jcim.3c01153.

(51) Mark, P.; Nilsson, L. Structure and Dynamics of the TIP3P, SPC, and SPC/E Water Models at 298 K. Journal of Physical Chemistry A 2001, 105 (43), 9954–9960. 10.1021/jp003020w.

(52) Maier, J. A.; Martinez, C.; Kasavajhala, K.; Wickstrom, L.; Hauser, K. E.; Simmerling, C. Ff14SB: Improving the Accuracy of Protein Side Chain and Backbone Parameters from Ff99SB. J. Chem. Theory Comput. 2015, 11 (8), 3696–3713. 10.1021/acs.jctc.5b00255.

(53) Ivani, I.; Dans, P. D.; Noy, A.; Pérez, A.; Faustino, I.; Hospital, A.; Walther, J.; Andrio, P.; Goñi, R.; Balaceanu, A.; Portella, G.; Battistini, F.; Gelpí, J. L.; González, C.; Vendruscolo, M.; Laughton, C. A.; Harris, S. A.; Case, D. A.; Orozco, M. Parmbsc1: A Refined Force Field for DNA Simulations. Nat. Methods 2015, 13 (1), 55–58. 10.1038/nmeth.3658.

(54) Wang, J.; Wang, W.; Kollman, P. A.; Case, D. A. Automatic Atom Type and Bond Type Perception in Molecular Mechanical Calculations. J. Mol. Graph. Model. 2006, 25 (2), 247–260. 10.1016/j.jmgm.2005.12.005.

(55) Case, D. A.; Cerutti, D. S.; Cruzeiro, V. W. D.; Darden, T. A.; Duke, R. E.; Ghazimirsaeed, M.; Giambaşu, G. M.; Giese, T. J.; Götz, A. W.; Harris, J. A.; Kasavajhala, K.; Lee, T.-S.; Li, Z.; Lin, C.; Liu, J.; Miao, Y.; Salomon-Ferrrer, R.; Shen, J.; Snyder, R.; Swails, J.; Walker, R. C.; Wang, J.; Wu, X.; Zeng, J.; Cheatham III, T. E.; Roe, D. R.; Roitberg, A.; Simmerling, C.; York, D. M.; Nagan, M. C.; Merz, K. M. Recent Developments in Amber Biomolecular Simulations. J. Chem. Inf. Model. 2025, 65 (15), 7835–7843. 10.1021/acs.jcim.5c01063.

(56) Hopkins, C. W.; Le Grand, S.; Walker, R. C.; Roitberg, A. E. Long-Time-Step Molecular Dynamics through Hydrogen Mass Repartitioning. J. Chem. Theory Comput. 2015, 11 (4), 1864–1874. 10.1021/ct5010406.

(57) Miyamoto, S.; Kollman, P. A. Settle: An Analytical Version of the SHAKE and RATTLE Algorithm for Rigid Water Models. J. Comput. Chem. 1992, 13 (8), 952–962. 10.1002/jcc.540130805.

(58) Davidchack, R. L.; Handel, R.; Tretyakov, M. V. Langevin Thermostat for Rigid Body Dynamics. Journal of Chemical Physics 2009, 130 (23), 234101. 10.1063/1.3149788.

(59) Feller, S. E.; Zhang, Y.; Pastor, R. W.; Brooks, B. R. Constant Pressure Molecular Dynamics Simulation: The Langevin Piston Method. J. Chem. Phys. 1995, 103 (11), 4613–4621. 10.1063/1.470648.

(60) Humphrey, W.; Dalke, A.; Schulten, K. VMD: Visual Molecular Dynamics. J. Mol. Graph. 1996, 14 (1), 33–38. 10.1016/0263-7855(96)00018-5.

(61) PTRAJ and CPPTRAJ: Software for Processing and Analysis of Molecular Dynamics Trajectory Data.

(62) Lu, X. J.; Olson, W. K. 3DNA: A Software Package for the Analysis, Rebuilding and Visualization of Three-Dimensional Nucleic Acid Structures. Nucleic Acids Res. 2003, 31 (17), 5108–5121. 10.1093/nar/gkg680.

(63) El Hassan, M. A.; Calladine, C. R. Two Distinct Modes of Protein-Induced Bending in DNA 1 1Edited by J. Karn. J. Mol. Biol. 1998, 282 (2), 331–343. 10.1006/jmbi.1998.1994.

