## Supplementary Information for "The Conformation of the Complementary Strand and the Deformation of the DNA Groove upon DDB2 Binding Justifies the Different Repair Rates for Cyclobutane Pyrimidine Dimers"

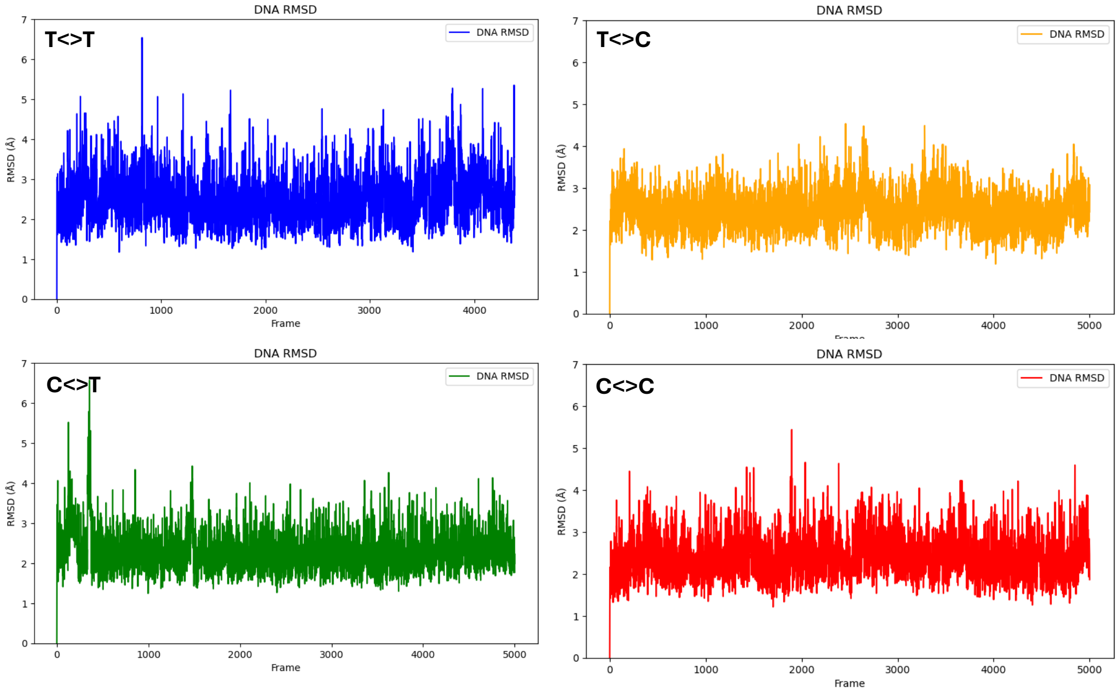


Figure S1. Time evolution of the Root Mean Square (RMSD) for the DNA only system.

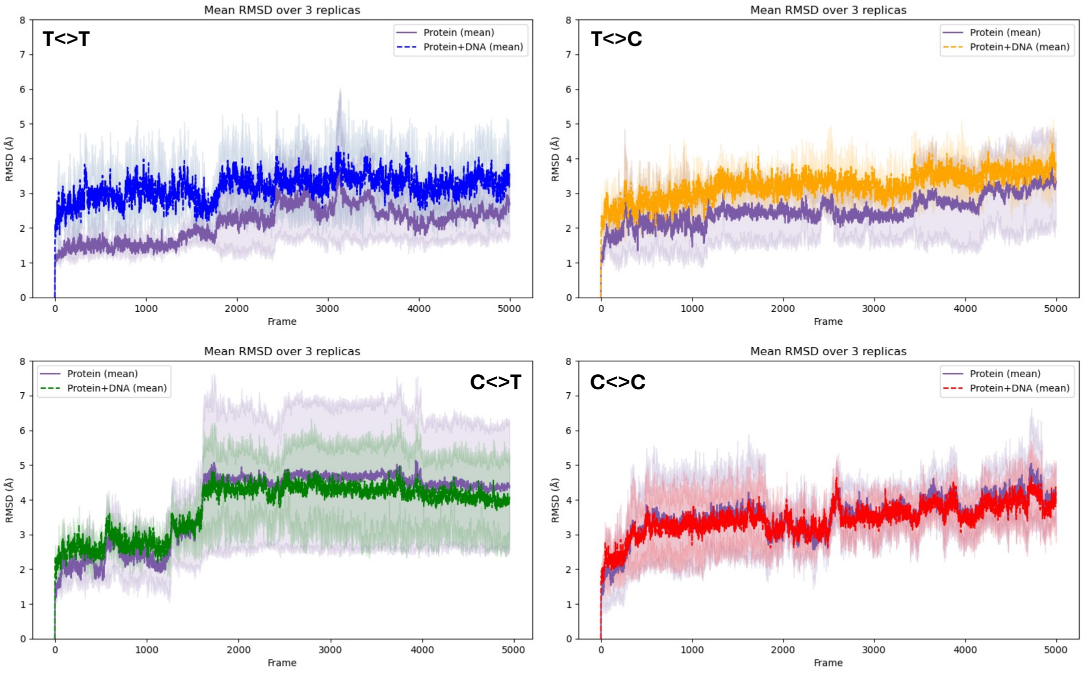


Figure S2. Time evolution of the Root Mean Square (RMSD) for the DNA/DDB2 system.

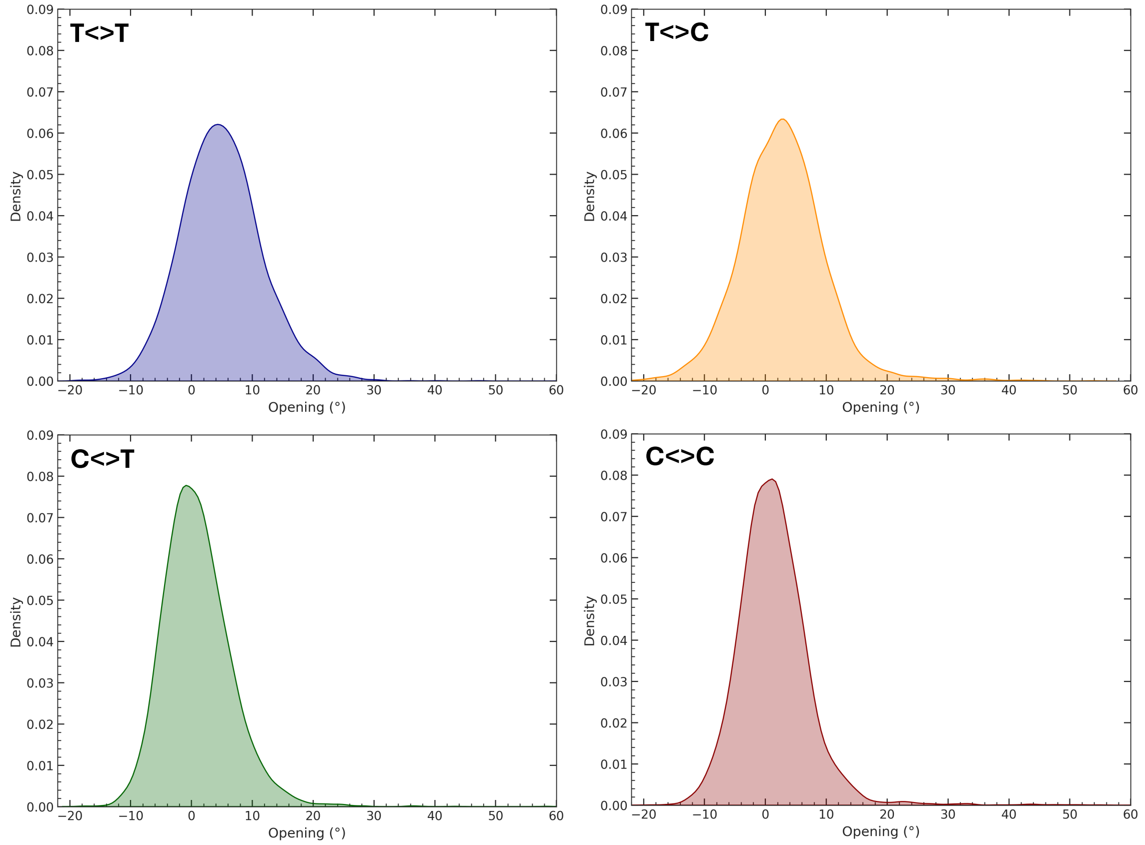


Figure S3. Distribution of the opening of the 5’ CPD base for naked DNA.

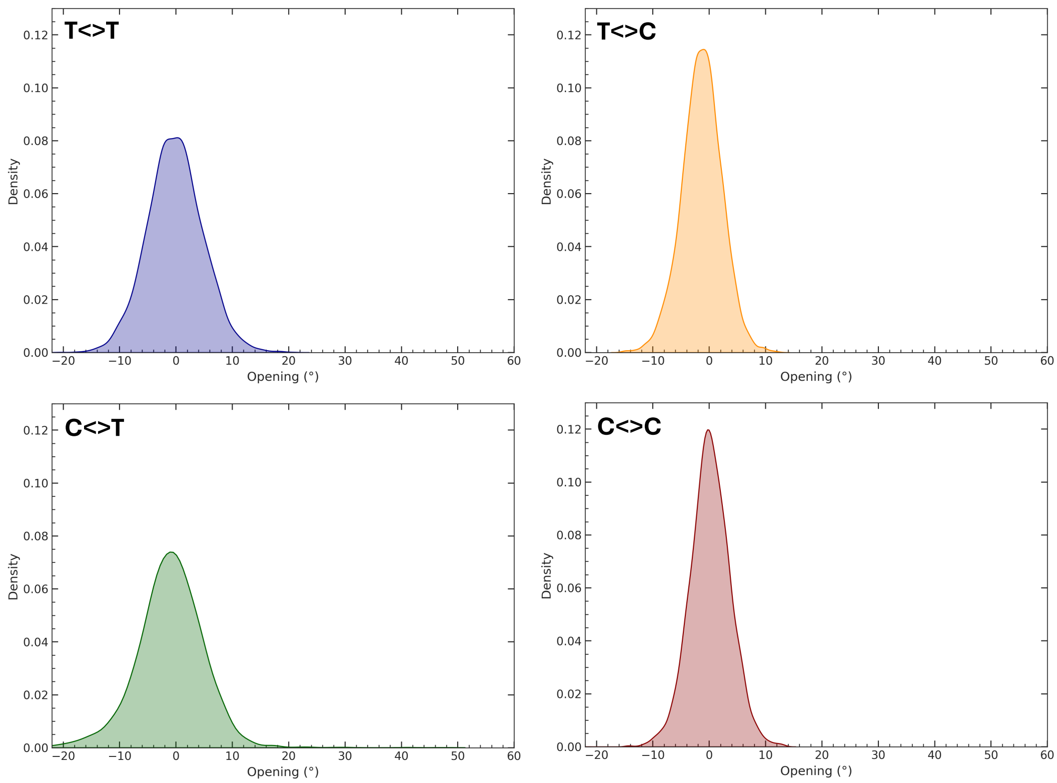


Figure S4. Distribution of the opening of the 3’ CPD base for naked DNA.

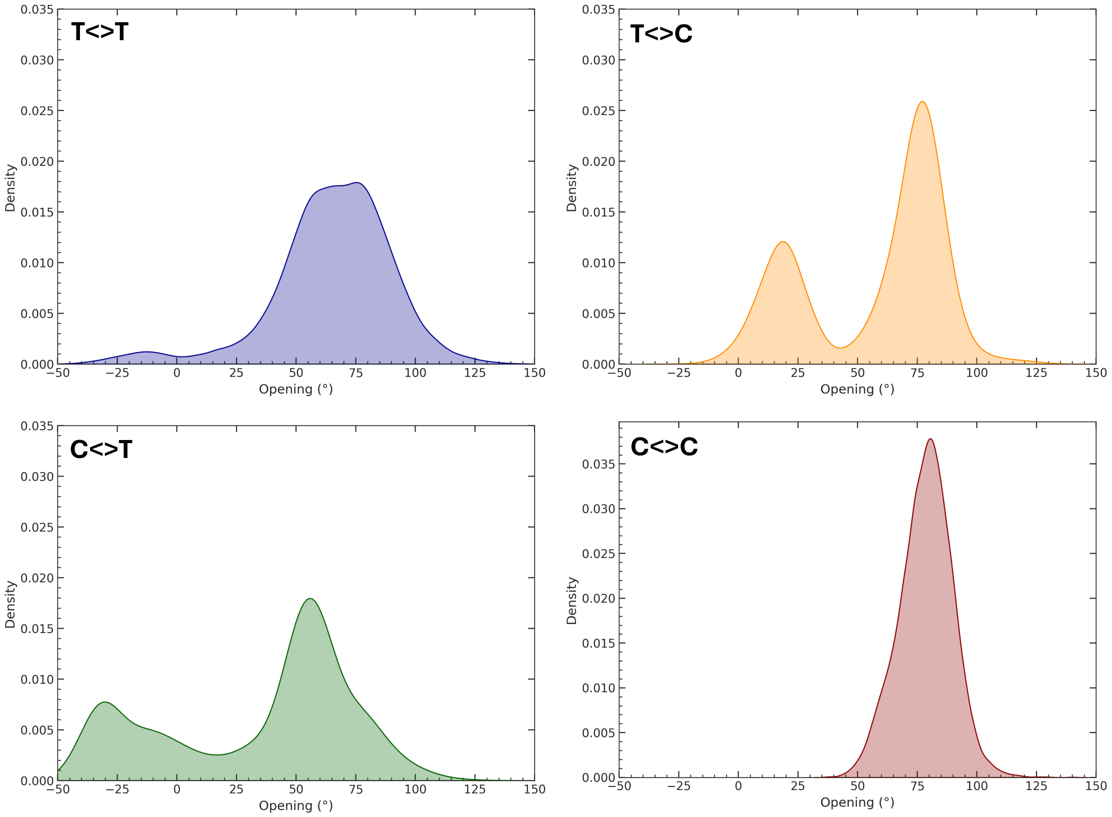


Figure S5. Distribution of Opening for the 3' damaged nucleotide on the ensemble of the DNA/DDB2 trajectories.

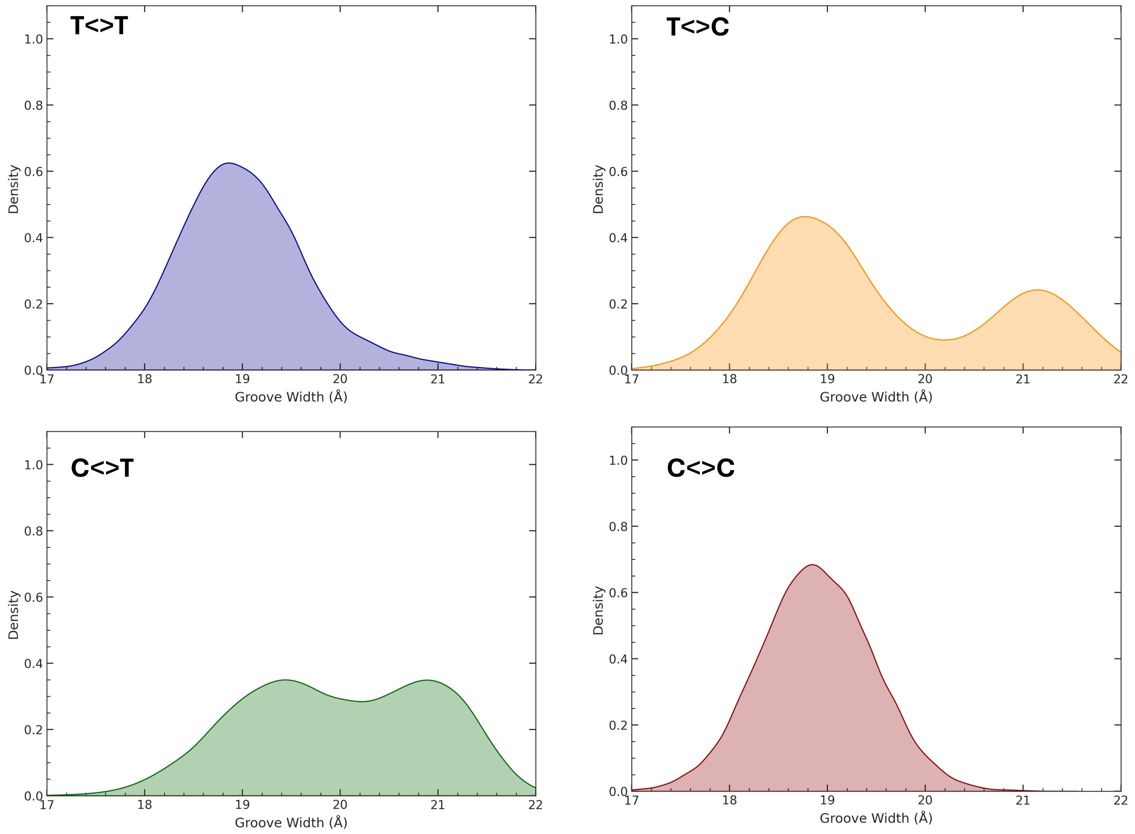


Figure S6. Distribution of the major groove width as extracted from the MD trajectory at the position +1 with respect to the CPD lesion on the ensemble of the DNA/DDB2 trajectories.

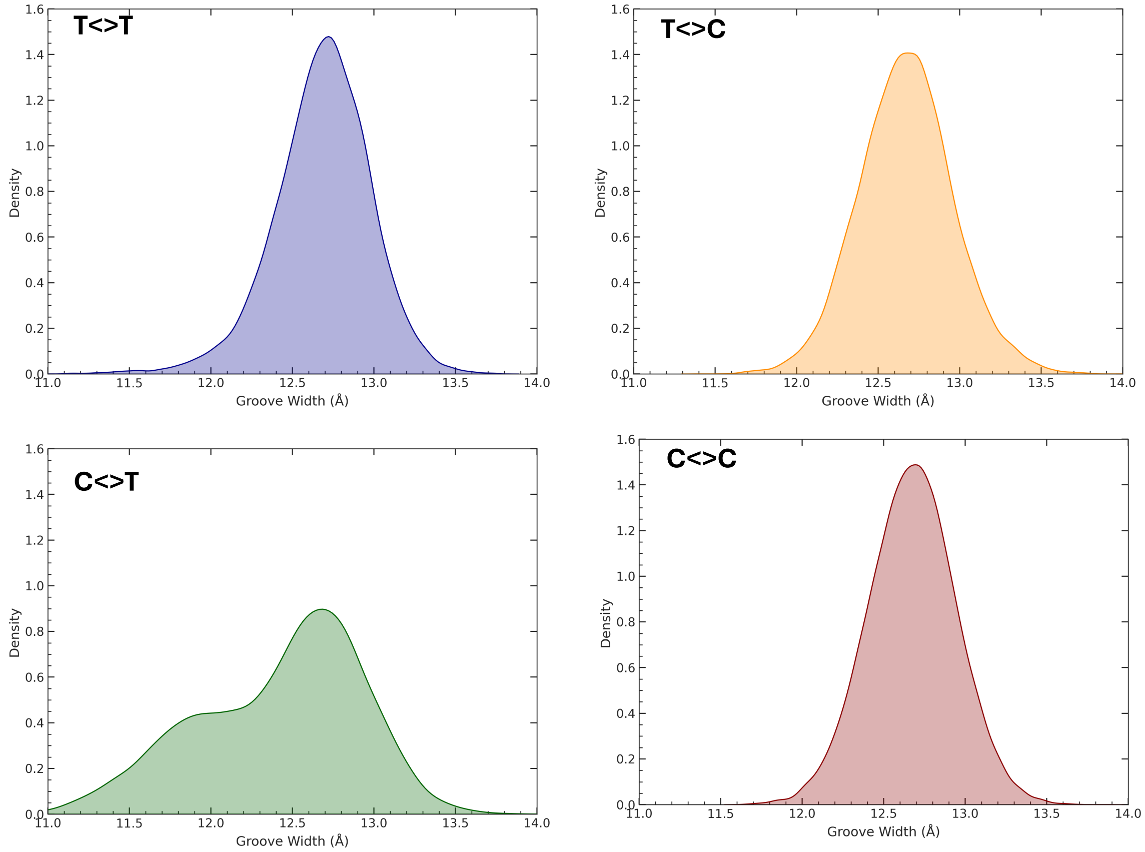


Figure S7. Distribution of the minor groove width as extracted from the MD trajectory at the position -1 with respect to the CPD lesion on the ensemble of the DNA/DDB2 trajectories.

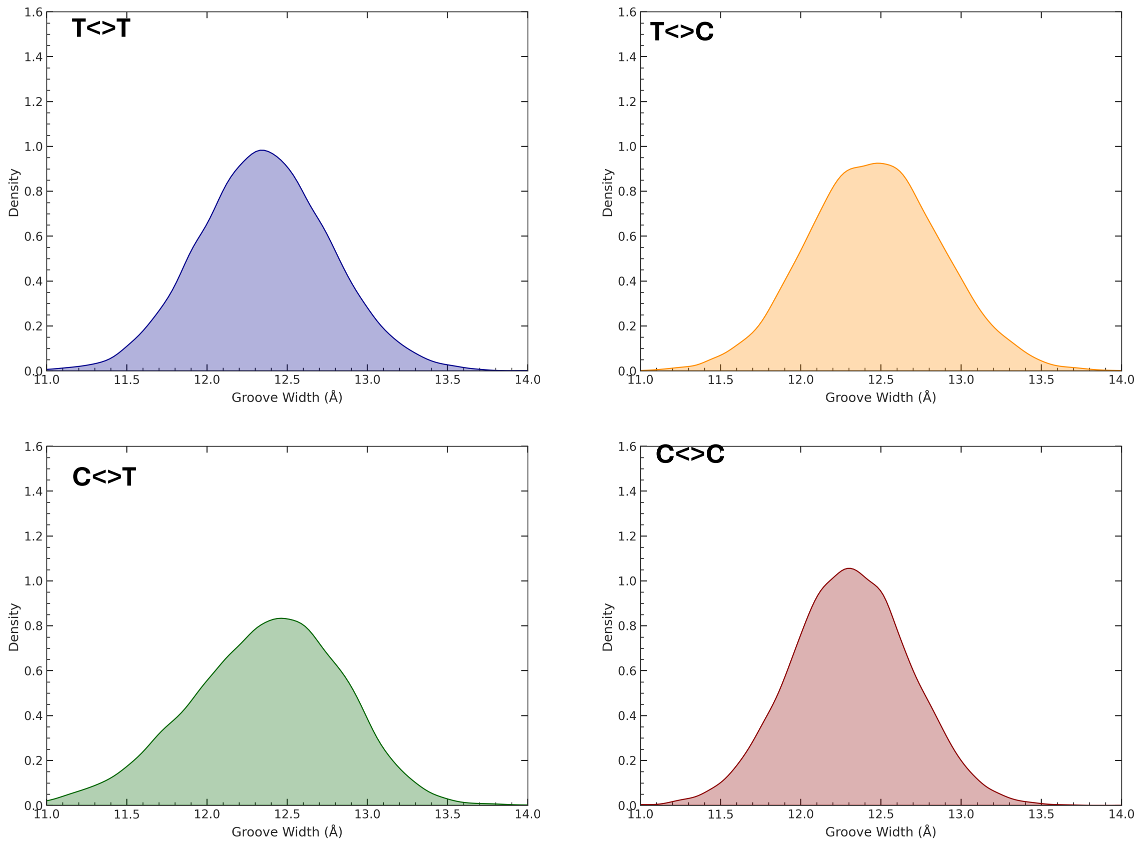


Figure S8. Distribution of the minor groove width as extracted from the MD trajectory at the position +1 with respect to the CPD lesion on the ensemble of the DNA/DDB2 trajectories.

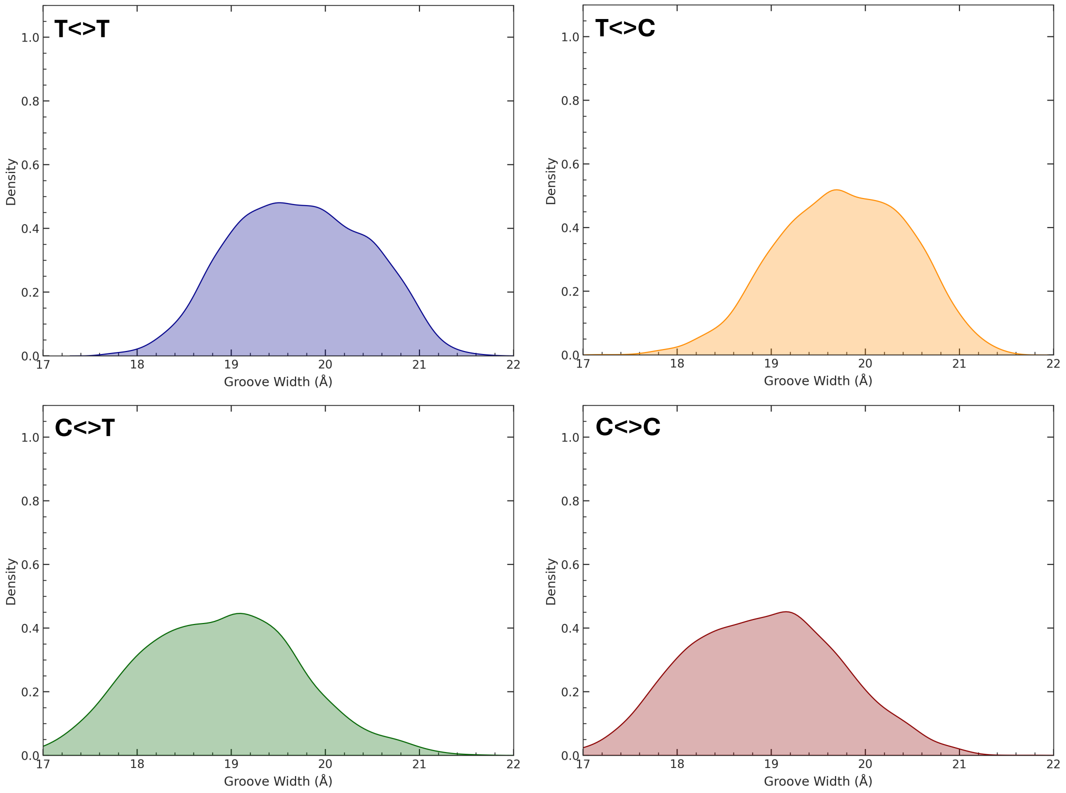


Figure S9. Distribution of the major groove width as extracted from the MD trajectory at the position -1 with respect to the CPD lesion on the naked DNA MD.

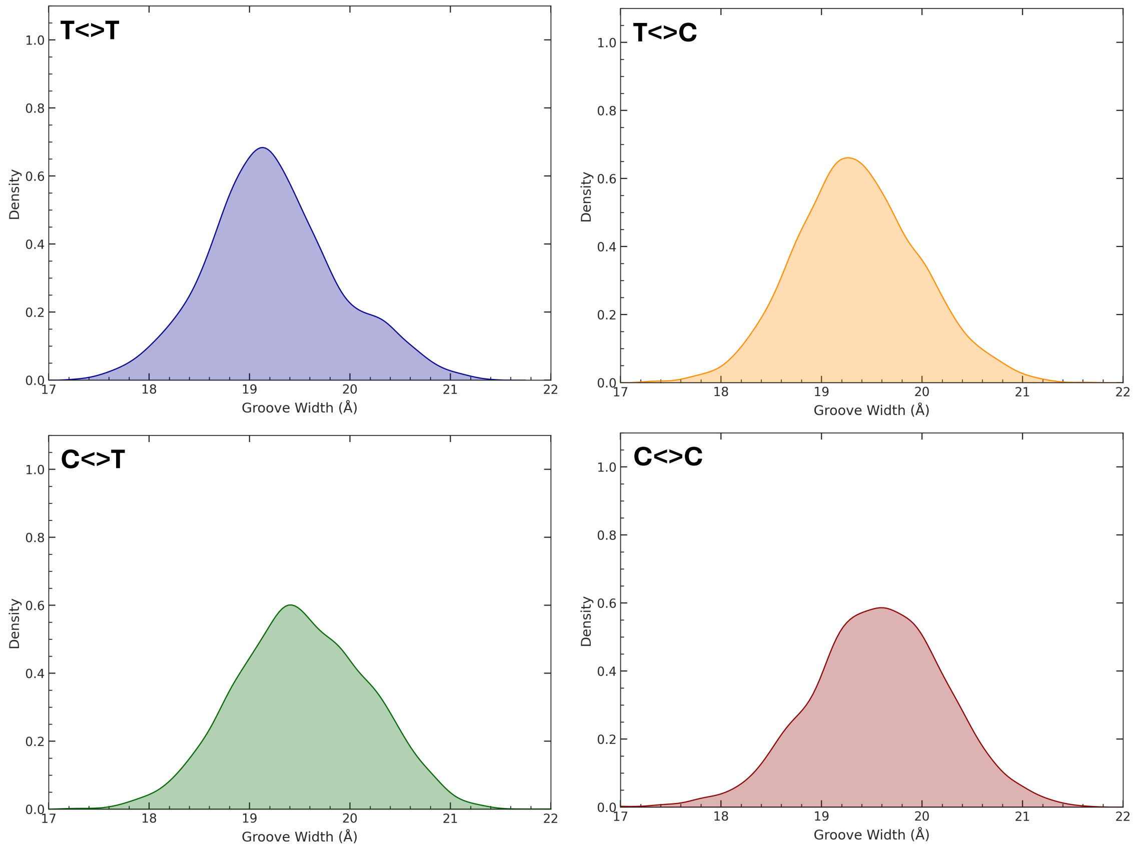


Figure S10. Distribution of the major groove width as extracted from the MD trajectory at the position +1 with respect to the CPD lesion on the naked DNA MD.

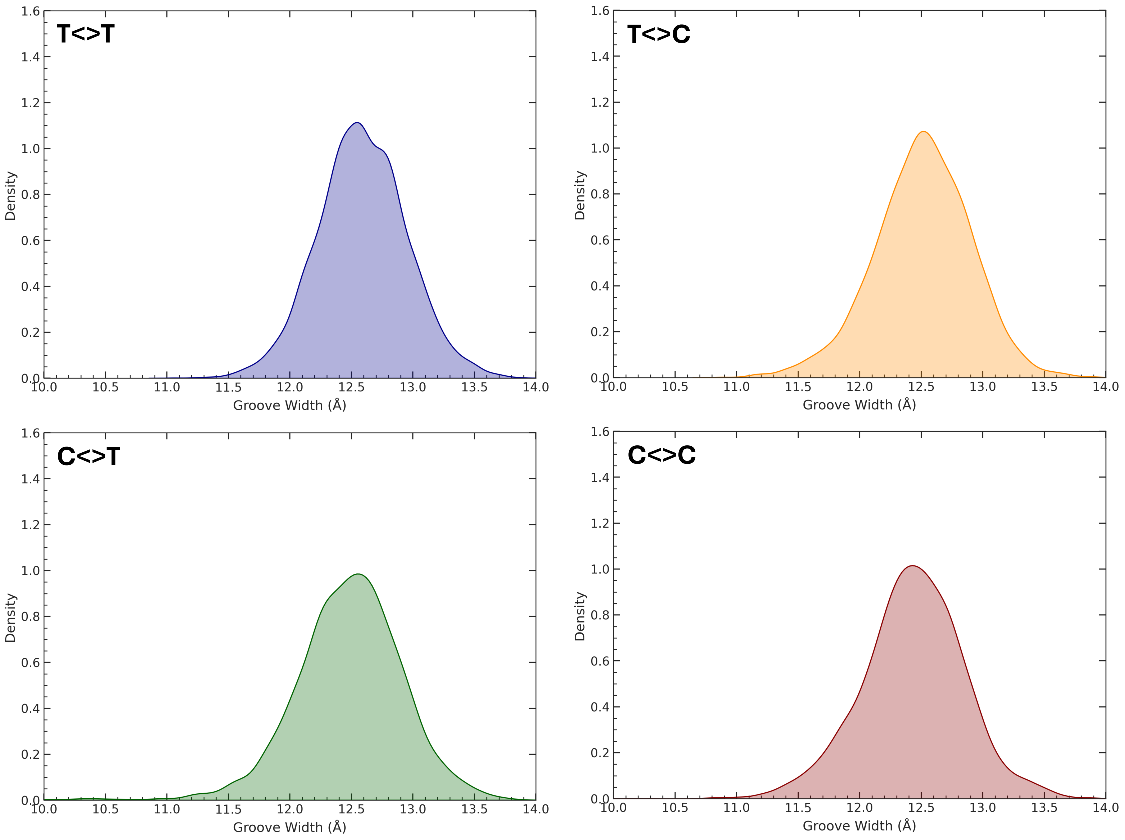


Figure S11. Distribution of the minor groove width as extracted from the MD trajectory at the position -1 with respect to the CPD lesion on the naked DNA MD.

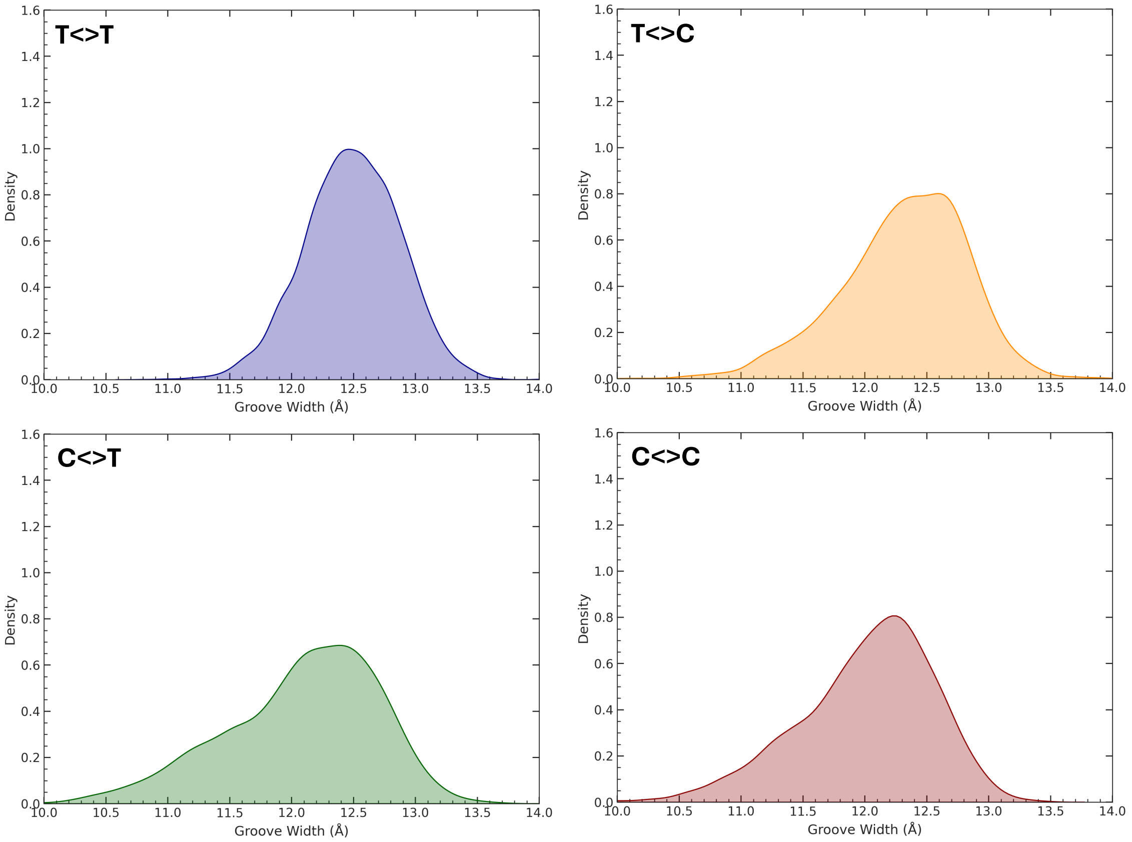


Figure S12. Distribution of the minor groove width as extracted from the MD trajectory at the position +1 with respect to the CPD lesion on the naked DNA MD.

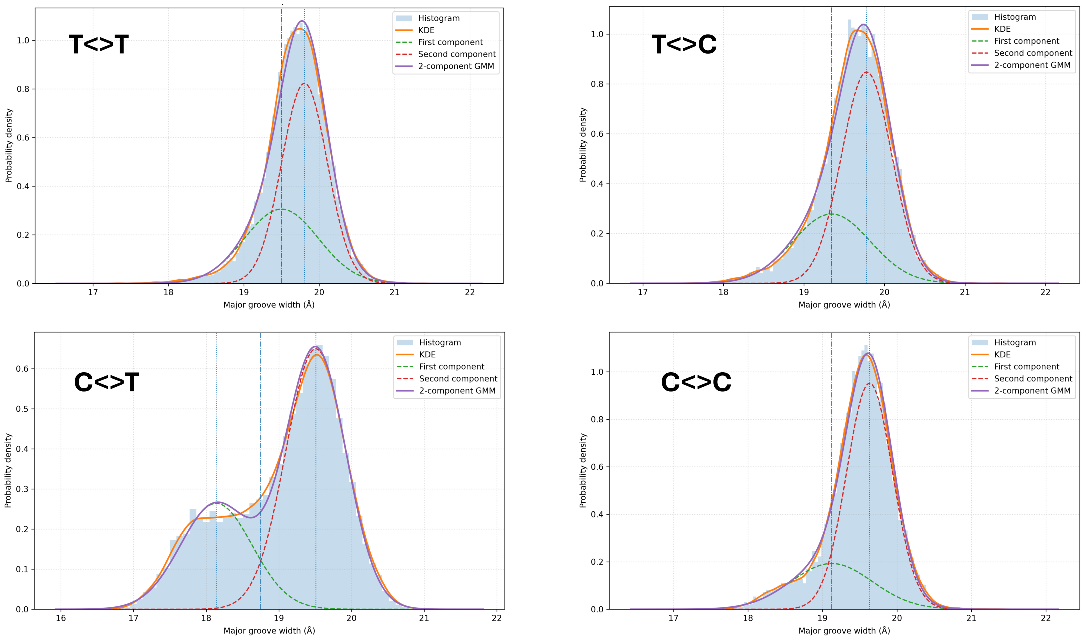


Figure S13. Gaussian Mixture Model (GMM) of the distribution of the major groove width at the -1 position.

| | | Mean (Å) | FWHM (Å) | Weight |
|---|---|---|---|---|
| T<>T | | | | |
| | First Component | 19.5 | 1.1 | 37.5% |
| | Second Component | 19.8 | 0.7 | 62.5% |
| T<>C | | | | |
| | First Component | 19.3 | 1.1 | 33.5% |
| | Second Component | 19.8 | 0.7 | 66.5% |
| C<>T | | | | |
| | First Component | 18.1 | 1.2 | 32.6% |
| | Second Component | 19.5 | 0.9 | 67.4% |
| C<>C | | | | |
| | First Component | 19.1 | 1.3 | 26.3% |
| | Second Component | 19.6 | 0.7 | 73.7% |

Table S1. Parameters of the two components identified by the GMM analysis to represent the distribution of the major groove width of the -1 position. The mean value, the full width at half maximum (FWHM) and the weight are given for each component.

| T<>T CPD | | | C<>T CPD | | |
|---|---|---|---|---|---|
| Name | Atom type | Charge | Name | Atom type | Charge |
| P | P | 1.1659 | P | P | 1.1659 |
| O1P | O2 | -0.7761 | O1P | O2 | -0.7761 |
| O2P | O2 | -0.7761 | O2P | O2 | -0.7761 |
| O5’ | OS | -0.4954 | O5’ | OS | -0.4954 |
| C5’ | CI | -0.0069 | C5’ | CI | -0.0069 |
| H5’ | H1 | 0.0754 | H5’ | H1 | 0.0754 |
| H5” | H1 | 0.0754 | H5” | H1 | 0.0754 |
| C4’ | CT | 0.1629 | C4’ | CT | 0.1629 |
| H4’ | H1 | 0.1176 | H4’ | H1 | 0.1176 |
| O4’ | OS | -0.3691 | O4’ | OS | -0.3691 |
| C1’ | CT | 0.0680 | C1’ | CT | 0.0680 |
| N1 | N* | -0.2491 | N1 | N* | -0.5378 |
| C6 | CT | 0.1238 | C6 | CT | 0.2118 |
| H6 | H1 | 0.0777 | H6 | H1 | 0.1404 |
| C5 | CT | 0.1877 | C5 | CT | -0.3621 |
| C7 | CT | -0.4262 | H5 | HC | 0.1288 |
| H71 | HC | 0.1282 | C4 | CM | 0.9422 |
| H72 | HC | 0.1282 | N4 | NT | -1.0819 |
| H73 | HC | 0.1282 | H41 | H | 0.4158 |
| C4 | C | 0.7381 | H42 | H | 0.4158 |
| O4 | O | -0.6280 | N3 | NC | -0.9224 |
| N3 | N | -0.6703 | C2’ | C | 1.0671 |
| H3 | H | 0.3600 | O2 | O | -0.6872 |
| C2 | C | 0.7883 | H1’ | H2 | 0.1963 |
| O2 | O | - 0.8283 | C3’ | CT | 0.0713 |
| H1’ | H2 | 0.1804 | H3’ | H1 | 0.0985 |
| C3’ | CE | 0.0713 | C2’ | CT | -0.0854 |
| H3’ | H1 | 0.0985 | H2’ | HC | 0.0718 |
| C2’ | CT | -0.0854 | H2” | HC | 0.0718 |
| H2’ | HC | 0.0718 | | | |
| H2” | HC | 0.0718 | | | |
| O3’ | OS | -0.5232 | O3’ | OS | -0.5232 |
| P | P | 1.1659 | P | P | 1.1659 |
| OP1 | O2 | -0.7761 | OP1 | O2 | -0.7761 |
| OP2 | O2 | -0.7761 | OP2 | O2 | -0.7761 |
| O5’ | OS | -0.4954 | O5’ | OS | -0.4954 |
| C5’ | C1 | -0.0069 | C5’ | C1 | -0.0069 |
| H5’ | H1 | 0.0754 | H5’ | H1 | 0.0754 |
| H5” | H1 | 0.0754 | H5” | H1 | 0.0754 |
| C4’ | CT | 0.1629 | C4’ | CT | 0.1629 |
| H4’ | H1 | 0.1176 | H4’ | H1 | 0.1176 |
| O4’ | OS | -0.3691 | O4’ | OS | -0.3691 |
| C1’ | CT | 0.0680 | C1’ | CT | 0.0680 |
| N8 | N* | -0.3265 | N1 | N | -0.2886 |
| C13 | CT | 0.0471 | C6 | CT | -0.1276 |
| H13 | H1 | 0.0913 | H6 | H1 | 0.1082 |
| C12 | CT | 0.0286 | C5 | CT | 0.1468 |
| C14 | CT | -0.4296 | C7 | CT | -0.2917 |
| H141 | HC | 0.1426 | H71 | HC | 0.0708 |
| H142 | HC | 0.1426 | H72 | HC | 0.0708 |
| H143 | HC | 0.1426 | H73 | HC | 0.0708 |
| C11 | C | 0.7270 | C4 | C | 0.7317 |
| O11 | O | -0.6015 | O4 | O | -0.6597 |
| N10 | N | -0.6838 | N3 | N | -0.8133 |
| H10 | H | 0.3853 | H3 | H | 0.3883 |
| C9 | C | 0.7252 | C2 | C | 0.8316 |
| O9 | O | -0.6217 | O2 | O | -0.6516 |
| H1’ | H2 | 0.1804 | H1’ | H2 | 0.1804 |
| C3’ | CE | 0.0713 | C3’ | CT | 0.0713 |
| H3’ | H1 | 0.0985 | H3’ | H1 | 0.0985 |
| C2’ | CT | -0.0854 | C2’ | CT | -0.0854 |
| H2’ | HC | 0.0718 | H2’ | HC | 0.0718 |
| H2” | HC | 0.0718 | H2” | HC | 0.0718 |
| O3’ | OS | -0.5232 | O3’ | OS | -0.5232 |

| T<>C CPD | | | C<>C CPD | | |
|---|---|---|---|---|---|
| Name | Atom type | Charge | Name | Atom type | Charge |
| P | P | 1.1659 | P | P | 1.1659 |
| O1P | O2 | -0.7761 | O1P | O2 | -0.7761 |
| O2P | O2 | -0.7761 | O2P | O2 | -0.7761 |
| O5’ | OS | -0.4954 | O5’ | OS | -0.4954 |
| C5’ | CI | -0.0069 | C5’ | CI | -0.0069 |
| H5’ | H1 | 0.0754 | H5’ | H1 | 0.0754 |
| H5” | H1 | 0.0754 | H5” | H1 | 0.0754 |
| C4’ | CT | 0.1629 | C4’ | CT | 0.1629 |
| H4’ | H1 | 0.1176 | H4’ | H1 | 0.1176 |
| O4’ | OS | -0.3691 | O4’ | OS | -0.3691 |
| C1’ | CT | 0.0680 | C1’ | CT | -0.0116 |
| N1 | N* | -0.2450 | N1 | N* | -0.3031 |
| C6 | CT | -0.0149 | C6 | CT | -0.0324 |
| H6 | H1 | 0.1976 | H6 | H1 | 0.2427 |
| C5 | CT | 0.0321 | C5 | CT | -0.2300 |
| C7 | CT | -0.3917 | H5 | HC | 0.1641 |
| H71 | HC | 0.1111 | C4 | CM | 0.9264 |
| H72 | HC | 0.1111 | N4 | NT | -1.1025 |
| H73 | HC | 0.1111 | H41 | H | 0.4583 |
| C4 | C | 0.7644 | H42 | H | 0.4583 |
| O4 | O | -0.6619 | N3 | NC | -0.8696 |
| N3 | N | -0.7666 | C2’ | C | 1.0325 |
| H3 | H | 0.3764 | O2 | O | -0.6658 |
| C2 | C | 0.8096 | H1’ | H2 | 0.1963 |
| O2 | O | -0.6612 | C3’ | CT | 0.0713 |
| H1’ | H2 | 0.1804 | H3’ | H1 | 0.0985 |
| C3’ | CE | 0.0713 | C2’ | CT | -0.0854 |
| H3’ | H1 | 0.0985 | H2’ | HC | 0.0718 |
| C2’ | CT | -0.0854 | H2” | HC | 0.0718 |
| H2’ | HC | 0.0718 | | | |
| H2” | HC | 0.0718 | | | |
| O3’ | OS | -0.5232 | O3’ | OS | -0.5232 |
| P | P | 1.1659 | P | P | 1.1659 |
| OP1 | O2 | -0.7761 | OP1 | O2 | -0.7761 |
| OP2 | O2 | -0.7761 | OP2 | O2 | -0.7761 |
| O5’ | OS | -0.4954 | O5’ | OS | -0.4954 |
| C5’ | C1 | -0.0069 | C5’ | C1 | -0.0069 |
| H5’ | H1 | 0.0754 | H5’ | H1 | 0.0754 |
| H5” | H1 | 0.0754 | H5” | H1 | 0.0754 |
| C4’ | CT | 0.1629 | C4’ | CT | 0.1629 |
| H4’ | H1 | 0.1176 | H4’ | H1 | 0.1176 |
| O4’ | OS | -0.3691 | O4’ | OS | -0.3691 |
| C1’ | CT | 0.0680 | C1’ | CT | -0.0116 |
| N1 | N* | -0.4945 | N1 | N | -0.4456 |
| C6 | CT | -0.0104 | C6 | CT | -0.0903 |
| H6 | H1 | 0.1354 | H6 | H1 | 0.1432 |
| C5 | CT | -0.4094 | C5 | CT | -0.0681 |
| H5 | HC | 0.1404 | H5 | HC | 0.1036 |
| C4 | CM | 1.0266 | C4 | CM | 0.8114 |
| N4 | NT | -1.1030 | N4 | NT | -1.0558 |
| H41 | H | 0.4241 | H41 | H | 0.4450 |
| H42 | H | 0.4241 | H42 | H | 0.4450 |
| N3 | NC | -0.9200 | N3 | NC | -0.8316 |
| C2 | C | 1.0216 | C2 | C | 1.0011 |
| O2 | O | -0.6902 | O2 | O | 0.6632 |
| H1’ | H2 | 0.1963 | H1’ | H2 | 0.1963 |
| C3’ | CT | 0.0713 | C3’ | CT | 0.0713 |
| H3’ | H1 | 0.0985 | H3’ | H1 | 0.0985 |
| C2’ | CT | -0.0854 | C2’ | CT | -0.0854 |
| H2’ | H1 | 0.0985 | H2’ | HC | 0.0718 |
| H2” | CT | -0.0854 | H2” | HC | 0.0718 |
| O3’ | OS | -0.5232 | O3’ | OS | -0.5232 |

Table S2. Atom types and RESP charges for the four CPD residue. For each photolesion, RESP charges were derived from the HF/6-31G* level of theory. They are reported in Table S2. We already derive these charges for the T<>T photolesion (1) and adopt the same procedure to compare the cytosine-containing photolesion on the same footing. The ground state geometry of each CPD has been previously optimized at B3LYP/6-31G* level.


1. Bignon,E., Gattuso,H., Morell,C., Dehez,F., Georgakilas,A.G., Monari,A. and Dumont,E. (2016) Correlation of bistranded clustered abasic DNA lesion processing with structural and dynamic DNA helix distortion. *Nucleic Acids Res.*, **44**, 8588-8599.
